# *Cauliflower Mosaic Virus* Utilizes Processing Bodies to Escape Translational Repression in Arabidopsis

**DOI:** 10.1101/2021.06.09.447751

**Authors:** Gesa Hoffmann, Amir Mahboubi, Damien Garcia, Johannes Hanson, Anders Hafrén

## Abstract

Viral infections impose extraordinary RNA stress on a cell, triggering cellular RNA surveillance pathways like RNA decapping, nonsense-mediated decay and RNA silencing. Viruses need to maneuver between these pathways to establish infection and succeed in producing high amounts of viral proteins. Processing bodies (PBs) are integral to RNA triage in eukaryotic cells with several distinct RNA quality control pathways converging for selective RNA regulation. In this study, we investigate the role of *Arabidopsis thaliana* PBs during *Cauliflower Mosaic Virus* (CaMV) infection. We find that several PB components are co-opted into viral replication factories and support virus multiplication. This pro-viral role was not associated with RNA decay pathways but instead, we could establish PB components as essential helpers in viral RNA translation. While CaMV is normally resilient to RNA silencing, PB dysfunctions expose the virus to this pathway, similar to previous observations on transgenes. Transgenes, however, undergo RNA Quality Control dependent RNA degradation, whereas CaMV RNA remains stable but becomes translationally repressed through decreased ribosome association, revealing a unique dependence between PBs, RNA silencing and translational repression. Together, our study shows that PB components are co-opted by the virus to maintain efficient translation, a mechanism not associated with canonical PB functions.

## INTRODUCTION

Eukaryotic gene expression is tightly regulated from RNA transcription to translation and decay. While transcription levels play an essential role in the central dogma, the importance of post-transcriptional control especially during stress-induced cellular reprogramming is becoming increasingly evident, as several studies revealed an extensive uncoupling between quantitative changes in transcriptomes and translatomes (Xu *et al*., 2017; Zid & O’Shea, 2014; Liu *et al*., 2013; Tebaldi *et al*., 2012; Branco-Price *et al*., 2005).

Due to the high energy cost and possible detrimental effects of uncontrolled protein translation, eukaryotic cells have evolved a network of pathways to govern and regulate mRNA translation, termed the “mRNA cycle” (Buchan & Parker, 2009). Here, cytoplasmic mRNAs are channeled between ribosomes and phase-separated cytoplasmic ribonucleo-protein (RNP) complexes, the RNA granules, in a triage between translation, non-translating storage, and degradation. Several kinds of constitutive and stress induced RNA granules have been identified and defined by their core protein constituents (Xing *et al*., 2020; Chantarachot & Bailey-Serres, 2018). The mRNA cycle involves two major types of cytoplasmic RNA granule termed processing bodies (PBs) and stress granules (SGs). RNAs are considered to shuffle between active translation at ribosomes and translationally repressed states at SGs (Buchan & Parker, 2009). In contrast, association with PBs was mainly associated with RNA degradation owing to the absence of translation initiation factors and to the highly conserved PB core components involved in RNA nonsense-mediated decay (NMD), miRNA-targeted gene silencing, deadenylation and decapping (Anderson & Kedersha, 2009). Yet, while PB proteins can facilitate translational repression (Xu & Chua, 2009), recent studies have shown that PB associated mRNAs can be stabilized and return to translation, expanding the multifunctionality of these RNA granules (Jang *et al*., 2019; Wang *et al*., 2018; Hubstenberger *et al*., 2017).

One hallmark of PBs is the accumulation of proteins involved in mRNA decapping. This process involves the removal of the 7-methyl-guanosine 5’-diphosphate (cap) and is essential for subsequent 5’- to 3’-end mRNA degradation. In *Arabidopsis thaliana* (hereafter referred to as Arabidopsis) decapping is carried out by the nudix hydrolase DECAPPING2 (DCP2) and its co- factors DCP1 and VARICOSE (VCS) (Xu *et al*., 2006). Several proteins function in decapping activation and PB assembly, including DCP5 and the SM-like (LSM) 1-7 complex (Perea-Resa *et al*., 2012; Xu & Chua, 2009). Uncapped RNAs are degraded by the cytoplasmic EXORIBONUCLEASE 4 (XRN4), which was also shown to accumulate in PBs (Yu *et al*., 2019; Souret *et al*., 2004). The decapping machinery is one part of the extensive RNA surveillance network present in PBs and it is tightly connected with and shares its targets with NMD (Chicois *et al*., 2018). NMD is governed by the surveillance factor UP FRAMESHIFT 1 (UPF1), which in combination with other factors monitors RNA integrity and marks aberrant RNAs for degradation. Both, the 5’-3’ degradation and NMD, together with other RNA quality control (RQC) components, ensure cellular RNA homeostasis during development and stress. Interestingly, UPF1 not only associates with PBs, but was also found to co-localize and shuffle between another class of cytoplasmic RNP granules, the small interfering (si)RNA bodies (Moreno *et al*., 2013). siRNA bodies are condensates of RNA DEPENDENT POLYMERASE6 (RDR6), SUPRESSOR OF GENE SILENCING3 (SGS3) and ARGONAUTE7 (AGO7), as well as other post-transcriptional gene silencing (PTGS) factors (Jouannet *et al*., 2012). These bodies can localize adjacent to PBs, and are proposed to store translationally repressed RNAs to triage them between PBs and RDR6-dependent PTGS, potentially through their interactions with UPF1 (Moreno *et al*., 2013; Jouannet *et al*., 2012).

Apart from their physical association, several connections and a tight inter-dependence of the RQC machinery and PTGS have been discovered in plants (Liu & Chen, 2016). An initial observation was the susceptibility of transgenes to suppression by RNA silencing in *dcp2* mutants (Thran *et al*., 2012). Subsequently, decapping mutants were found to accumulate novel classes of endogenous siRNAs that arose through the cytoplasmic RDR6 pathway (Martinez de Alba *et al*., 2015). In line with the central role of RDR6 in this process, its knock-out rescued the seedling lethality in the severe decapping mutants *vcs6* and *dcp2* (Martinez de Alba *et al*., 2015). The fact that major cytoplasmic RQC pathways and PTGS center around PBs, makes these RNA granules prime targets for virus resistance, but also for manipulation by viruses.

Viruses challenge the RQC and PTGS machineries through their massive production of RNAs during replication. Indeed, targeting of viral RNAs by RNA silencing is one of the major defense pathway plants employ against viruses and in turn, viruses have frequently evolved RNA silencing suppressors to overcome it (Csorba *et al*., 2015). The role of RQC and PBs during plant virus infections is currently not well understood, and how these pathways further co-operate with the RNA silencing defense and NMD against viruses is unknown. NMD can restrict viral accumulation while at the same time being a target of viral manipulation (May *et al*., 2020; Garcia *et al*., 2014). Several PB proteins were shown to regulate viral infections; Carbon Catabolite Repression 4 (CCR4) facilitates *Barley yellow striate mosaic virus* replication in barley (Zhang *et al*., 2020), VCS supports Potato virus A infection (De *et al*., 2020; Hafrén *et al*., 2015) and overexpression of several PB components was shown to restrict *Turnip Mosaic Virus (Li & Wang, 2018)*. It is becoming increasingly evident that RNA surveillance pathways, including RNA silencing and decapping, play a major role in translational regulation not only through degradation of aberrant RNAs, but by eliciting translational repression on endogenous targets (Hung & Slotkin, 2021; Iwakawa *et al*., 2020; Wu *et al*., 2020; Jang *et al*., 2019; Lanet *et al*., 2009; Xu & Chua, 2009; Brodersen *et al*., 2008). As part of antiviral translational regulation, different animal viruses are frequently challenged with a global shut-down of protein production. *In planta*, this has only been observed for geminiviruses (Zorzatto *et al*., 2015) and for example the plant virus *Cauliflower mosaic virus* (CaMV) increases global polysome-RNA association in infected cells (Park *et al*., 2001). Investigating to what extent plant viruses are subject to translational repression by RNA surveillance pathways, how this suppression is regulated, and which counterstrategies viruses have evolved is imperative for disease management and can lead to the discovery of novel functions in endogenous pathways, as previous research shows (Jaafar & Kieft, 2019; Stern-Ginossar *et al*., 2019; Pooggin & Ryabova, 2018; Miras *et al*., 2017).

In this study, we investigated the role of PBs and decapping components during viral infection with the pararetrovirus *Cauliflower mosaic virus* (CaMV; family *Caulimoviridae*) in the model plant Arabidopsis and provide an analysis of the interdependencies between three major RNA surveillance mechanisms, RNA decapping, NMD and RNA silencing, during viral infection. We show that at least three hallmark proteins of PBs are targeted to the viral factories (VFs) of CaMV and that these proteins are important for virus accumulation. We demonstrate that PBs serve a pro-viral role during CaMV infection by alleviating translational repression through RNA silencing. Our findings establish PBs as hubs for viral manipulation and as an escape route for plant viruses to evade RNA silencing.

## RESULTS

### PB components re-localize during CaMV infection

To visualize PB dynamics during CaMV infection, we used marker lines expressing GFP-tagged canonical PB proteins (DCP1-GFP, DCP5-GFP, LSM1a-GFP and GFP-VCS) (Chicois *et al*., 2018; Roux *et al*., 2015; Motomura *et al*., 2012). In mock conditions the markers showed a cytoplasmic distribution with varying degrees of condensation into droplet-like foci (Figure 1A, upper panel). LSM1a-GFP fusion protein accumulated evenly in the cytoplasm, with no visible PB assembly, while GFP-VCS and DCP5-GFP were present both in foci and soluble and DCP1- GFP mainly assembled in foci. These localization patterns were similar to those described previously (Chicois *et al*., 2018; Perea-Resa *et al*., 2016; Motomura *et al*., 2015; Roux *et al*., 2015). Before analyzing infection dynamics, we established how the markers behaved after heat shock (HS) application (Motomura *et al*., 2015). The number of detectable foci after HS increased drastically and was comparable for all markers, pointing towards a directed co-assembly during stress (Figure 1A,B). This is consistent with earlier findings that some PB proteins, including LSM1a, associate with PBs only upon stress (Guzikowski *et al*., 2019; Perea-Resa *et al*., 2016). Importantly, this analysis confirms functionality of the marker lines in our conditions.

**Figure 1:**
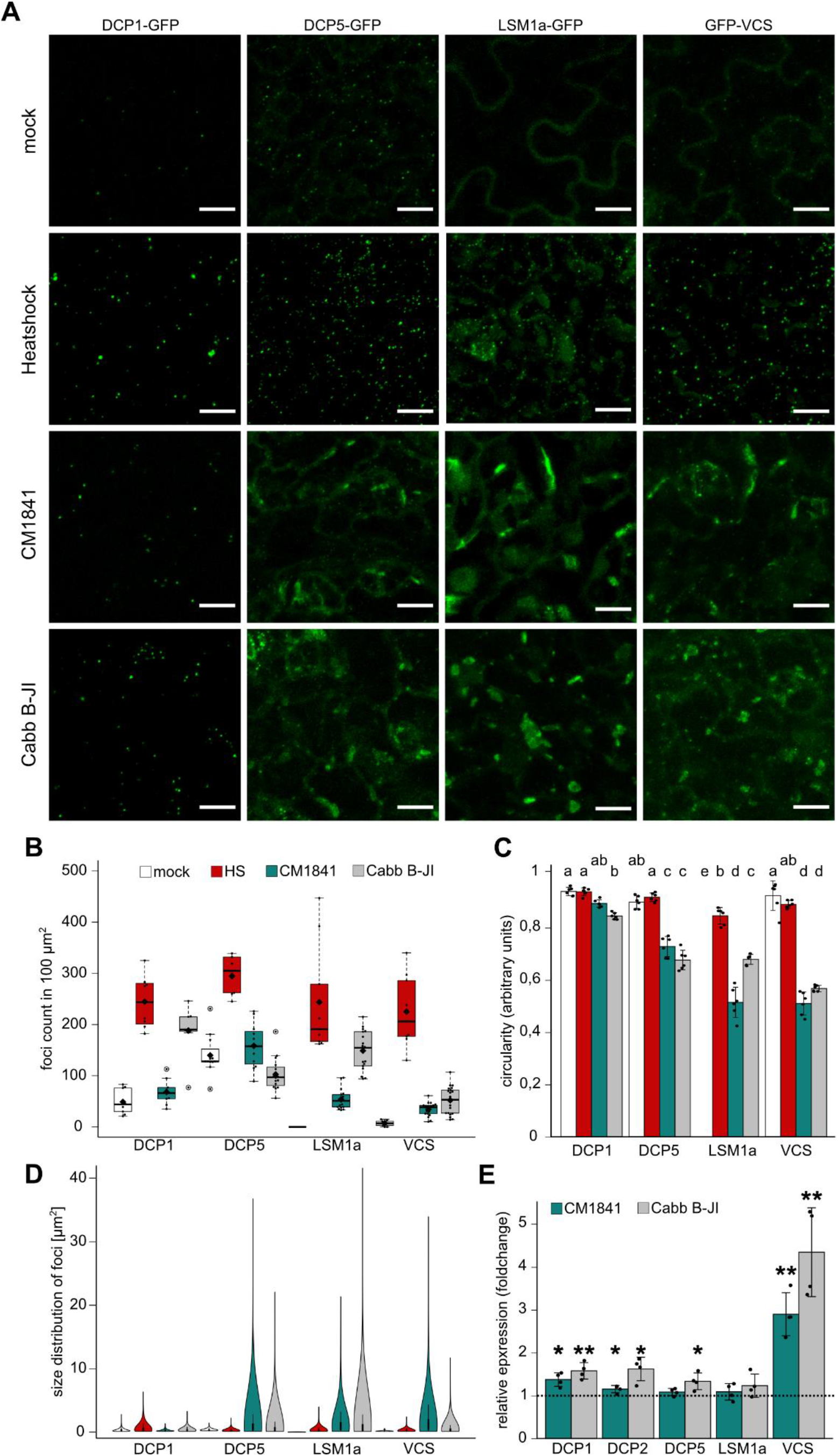
CaMV infection causes re-localization of PB proteins to novel structures. (A) Localization of four canonical PB markers in control conditions, after heat shock and 21 dpi with CaMV strains CM1841 and Cabb B-JI. The representative images are composed of confocal Z-stacks (Scale bars = 10 µm). (B) Count of fluorescent foci in 100 µm^2^ corresponding to treatments in (A). Counts were performed from randomly chosen areas using ImageJ and a custom pipeline. Solid lines represent median of the data, diamonds the average. (C) Average circularity of detected foci in each condition determined by ImageJ circularity masking. Significant differences were determined by one-way ANOVA coupled with Tuckey HSD test (alpha = 0.05), letters indicate statistical groups. Values calculated from six independent replicates. (D) Size distribution of detected foci corresponding to (B). (B/D): Values calculated from nine z-stacks of three plants for each replicate. All experiments were replicated at least three times independently. (E) Relative expression (foldchange) of PB components 21 dpi compared to mock (dashed line) (n=4). Experiment repeated three times independently. Statistical significance was determined by student-t test (*P < 0.05; **P < 0.01).

Upon infection with two CaMV strains (CM1841 and Cabb B-JI), the PB marker proteins formed two morphologically distinct classes of visible structures in systemic leaves 21 days post infection (dpi) (Figure 1A, lower panels). The number of DCP1-marked foci increased especially during Cabb B-JI infection without any apparent change in morphology (Figure 1A-D). LSM1, VCS and DCP5 markers also accumulated in small DCP1-like foci upon CaMV infection, but most striking was their prominent assembly into large, irregular shaped structures not seen with DCP1 (Figure 1A-D). The large structures were less abundant than the droplets for the three markers and had a distorted circularity which was not seen after HS or in the DCP1 marker (Figure 1C-D). We never detected comparable structures in either control conditions or after heat stress with any of the markers, while they were always found abundantly with both CaMV strains, with slight variation in numbers and size. Interestingly, these structures grew in size and decreased in numbers during the infection time course, indicating a fusion of them in infected cells (supplemental Figure 1). To validate the findings and confirm that the same structures were indeed marked by different PB markers, we established two double marker lines with GFP- VCS/DCP1-RFP and GFP-VCS/LSM1a-RFP. Both RFP-tagged markers showed the same localization patterns as the GFP-tagged constructs in mock and HS conditions. Interestingly, only a fraction of DCP1 and VCS co-localized in mock conditions, while co-assembly after HS confirmed the stress-dependent co-accumulation of PB markers (supplemental Figure S2). During CM1841 infection LSM1a-RFP and GFP-VCS both marked the same large, irregular structures, while DCP1-RFP localized in smaller foci adjacent to VCS structures (supplemental Figure S2). Thus, the double marker lines confirmed a co-assembly of three PB proteins in virus-induced structures, and an adjacent recruitment, but exclusion of DCP1 from them.

The increased GFP signal in virus infected plants, led us to test whether transcription of PB components was altered during infection. The transcript levels were elevated for DCP1 and more strongly for VCS with both CaMV strains but not for the other components (Figure 1E), showing that the dynamic shifts in foci counts and morphology are likely a re-distribution of previously soluble protein rather than transcription-driven *de novo* protein synthesis. In conclusion, CaMV infection causes condensation and a drastic re-localization of several PB components into large virus-induced structures.

### CaMV sequesters PB components into viral factories

The re-localization of LSM1, VCS and DCP5 into the morphologically distinct structures during CaMV infection, suggested that these could be virus induced inclusions. CaMV assembles two kinds of cytoplasmic inclusions; the spherical transmission bodies formed mainly by viral protein P2 and the more irregularly shaped VFs formed mainly by viral P6 (Espinoza *et al*., 1991; Martelli & Castellano, 1971). Heterologous co-expression of six CaMV proteins with PB proteins in *Nicotiana benthamiana* showed that viral P6 protein co-localized with DCP1, DCP5 and VCS (supplemental Figure S3), suggesting that PBs associate with VFs during CaMV infections. This prompted us to investigate co-localization of PB-markers with viral factories during CaMV infection. We used transgenic P6-mRFP expressing PB marker lines to investigate the association of DCP1, DCP5, LSM1 and VCS with VFs. Under control conditions, P6 is mostly soluble in the cytoplasm, with occasional foci formation (supplemental Figure 4). Some, but not all these foci were co-localizing with DCP1, DCP5 and VCS, indicating an association of the proteins already in the absence of infection (supplemental Figure 4, white arrows). During infection, the P6-mRFP protein assembled to mark the characteristic large VFs, which in addition accumulated DCP5, LSM1a and VCS. (Figure 2A). DCP1 foci accumulated around, but not within the VFs. Translation inhibition through trapping of ribosomes on mRNA by Cycloheximide (CHX) leads to the disassembly of canonical PBs (Motomura *et al*., 2015; Teixeira *et al*., 2005). In our conditions, CHX treatment of the DCP1-GFP and DCP5-GFP marker line after mock or CaMV infection, confirmed the dissociation of canonical PBs after CHX treatment, but the irregular VFs were still marked by DCP5 albeit signal intensity was lower in CHX treated samples (Figure 2B, lower panel). DCP1 bodies disappeared after treatment regardless of viral infection (Figure 2B, upper panel). This shows that DCP5 in VFs is dynamically less responsive to depletion of RNA supply from ribosomes than canonical PBs, possibly owing to VF size or other distinct physicochemical properties including interactions with the VF matrix.

**Figure 2:**
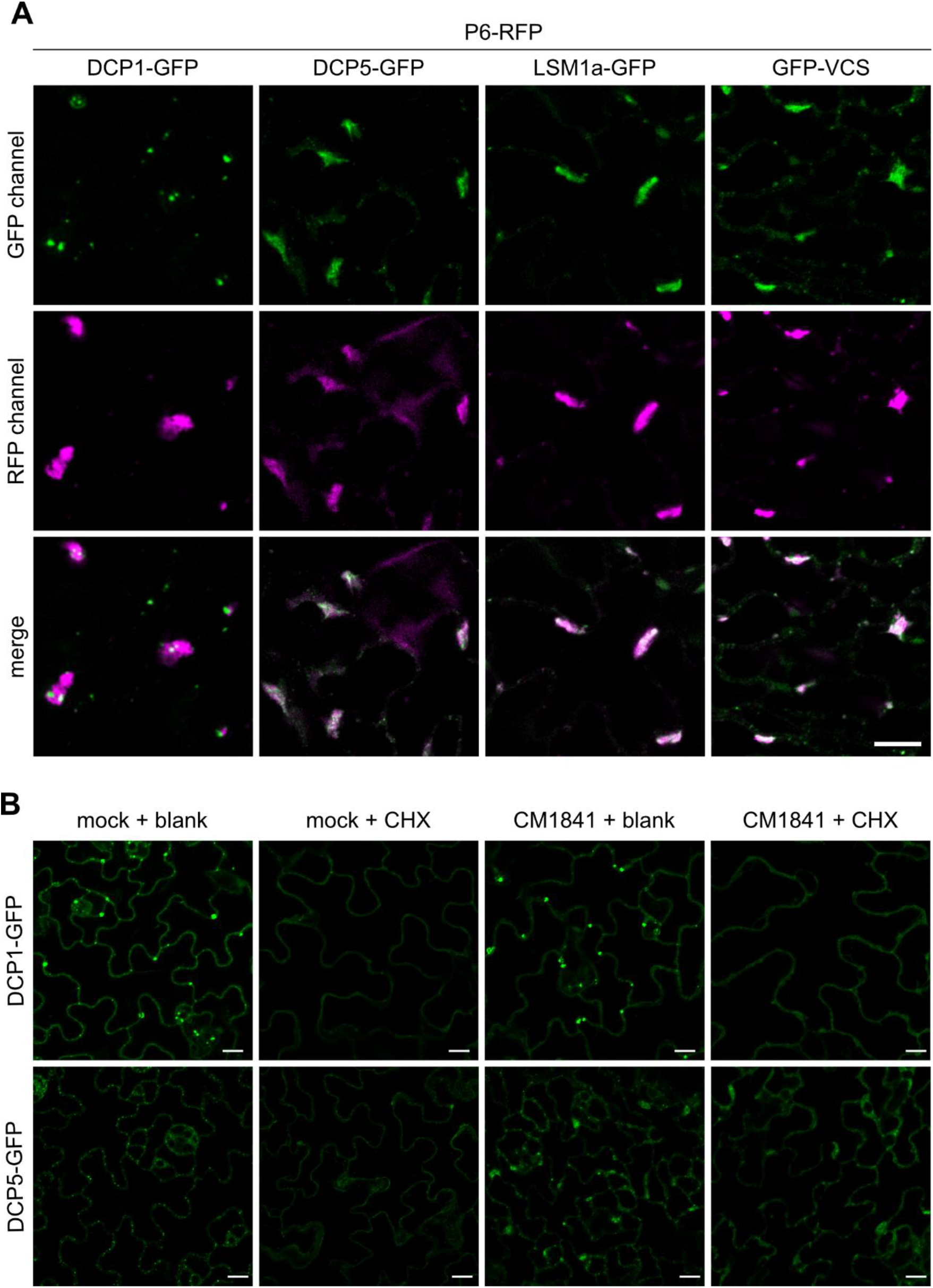
CaMV co-opts PB components into VFs. (A) Co-localization of P6-RFP with GFP-tagged PB markers in transgenic Arabidopsis 21 dpi with CaMV strain CM184I. Representative single plane images are shown (Scale bars = 10 µm). Experiments were replicated in independent transformants. (B) Distribution of DCP1-GFP and DCP5-GFP marker 21 days after mock or CaMV infection and 1h after 200 µM cycloheximide (CHX) or blank infiltration. Images represent single plane micrographs (Scale bars = 10 µm). DCP1- GFP were imaged with a higher exposure to ensure visualization of the soluble fraction.

### Disruption of PB functions attenuates CaMV infection

The VFs formed by CaMV P6 protein are electron dense, RNA and protein-rich structures with essential roles in the viral life cycle (Schoelz & Leisner, 2017; Martelli & Castellano, 1971). VFs are proposed to be sites of active viral RNA translation, reverse transcription, and packaging of viral genomic DNA in particles. Considering the re-localization of PB components to viral replication sites, we next investigated the role of PBs in CaMV disease by analyzing infection phenotypes in mutants disrupted in PB functions. The null mutant *lsm1a/b* (hereafter referred to as *lsm1*) and knock-down mutant *dcp5* were chosen for this study, because both mutations cause a reduction in PB formation and PB size, as well as an over-accumulation of capped mRNAs (Perea-Resa *et al*., 2016; Perea-Resa *et al*., 2012; Xu & Chua, 2009). Importantly, these mutants are not postembryonic lethal in contrast to null mutants of DCP1, DCP2 and VCS (Xu *et al*., 2006), and grow well enough for virus infection experiments. The *lsm1* and *dcp5* plants showed developmental phenotypes, including slightly delayed germination, mild dwarfism and leaf serrations (Figure 3A). Additionally, the null-mutant of the cytoplasmic exonuclease *xrn4* was used that is not impaired in PB biogenesis and mRNA decapping, but over-accumulates uncapped RNAs (Nagarajan *et al*., 2019). The *xrn4* plants were morphologically not distinguishable from Col-0 plants in our growth conditions.

**Figure 3:**
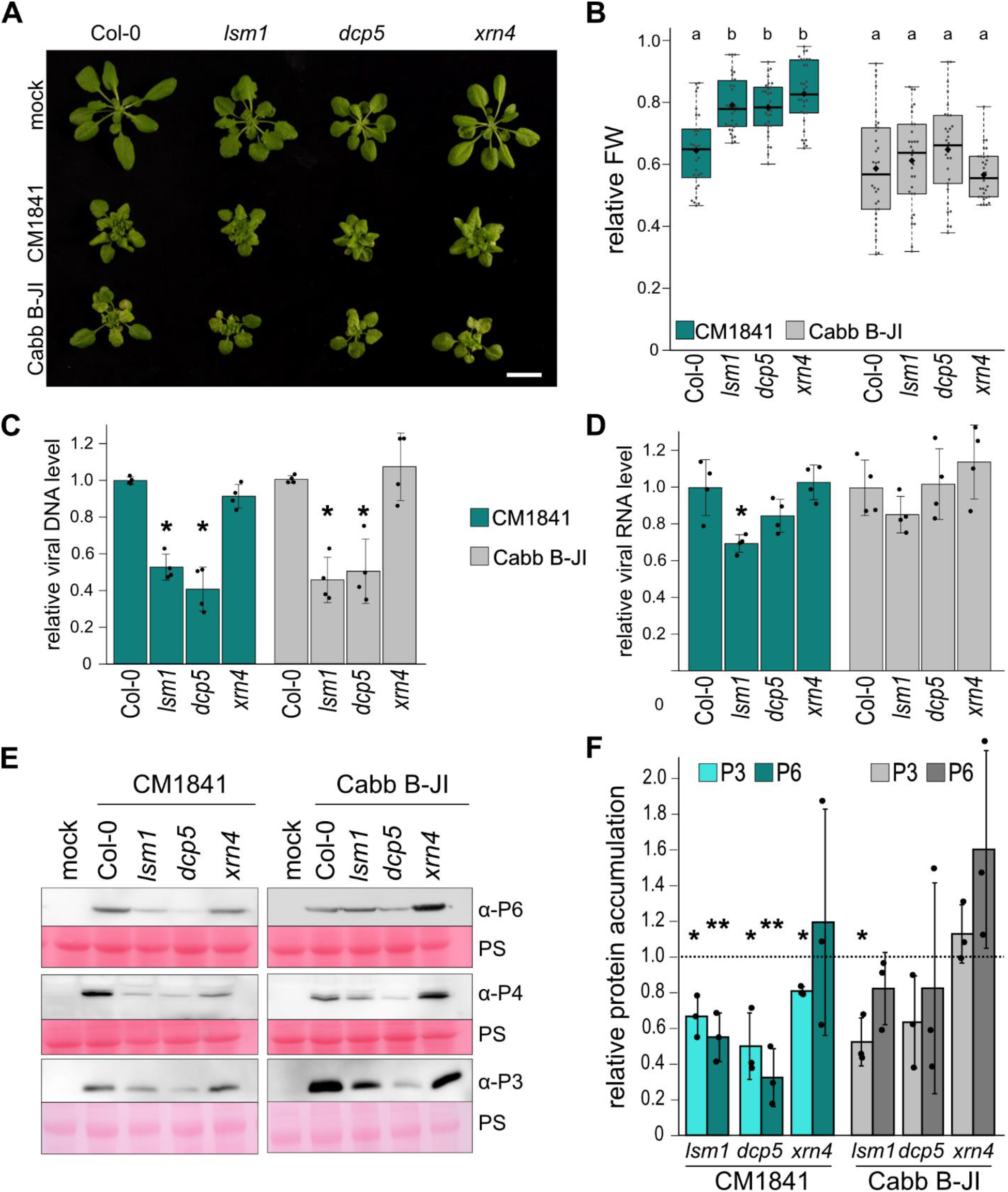
CaMV disease is attenuated in *lsm1* and *dcp5* mutants. **(A)** Virus-induced symptoms in Col-0, *lsm1*, *dcp5* and *xrn4* plants at 21 dpi. CM1841 and Cabb B-JI infected plants are compared to mock infected plants (Scale bar = 2 cm). (B) Relative fresh weight accumulation of CaMV infected compared to mock plants at 21 dpi (n = 30). Solid lines represent median of the data, diamonds the average. Statistical significance was determined by one-way ANOVA coupled with Tuckey HSD test (alpha = 0.05), letters indicate statistical groups. **(C)** Viral DNA accumulation in systemic leaves of Col-0 and mutant plants at 21 dpi, determined by qPCR. Values represent means ± SD (n = 4) relative to Col-0 plants and normalized to *18s* ribosomal DNA as the internal reference. **(D)** *35s* RNA levels of CaMV were determined by qRT-PCR in systemic leaves at 21 dpi. Values represent means ± SD (n = 4) relative to Col-0 plants and normalized to *PP2a*. **(E)** Immunoblot analysis of CaMV P3, P4 and P6 protein in Col-0, *lsm1*, *dcp5* and *xrn4* plants in systemic leaves. Total proteins were extracted at 21 dpi and probed with specific antibodies. Mock-infected plants were used as control for signal background. Ponceau S (PS) staining served as loading control. **(F)** Accumulation of CaMV P3 and P6 proteins in all genotypes in systemic leaves at 21 dpi quantified by direct ELISA. Values represent means ± SD (n = 3) in arbitrary units relative to Col-0 plants (dashed line). Statistical significance was determined by Student’s t test for C, D and F (*P < 0.05; **P < 0.01). All experiments **(A-F)** were repeated at least three times from independent infections.

Upon infection with CaMV, all mutants showed similar levels of stunting, vein bleaching, rosette distortion and leaf wrinkling compared to Col-0 (Figure 3A), with prominent symptoms appearing at 12 (Cabb B-JI) or 14 (CM1841) dpi. The relative fresh weight of CaMV infected compared to mock inoculated plants was taken as a measure for disease severity (Hafren *et al*., 2017). The fresh weight loss was less severe in all three mutants compared to Col-0 for the milder CM1841 strain, and unaltered for Cabb B-JI (Figure 3B). In general, Cabb B-JI infection caused stronger but also more variable infection phenotypes, possibly masking potential effects of PB disruption on fresh weight loss.

To establish viral load in the mutants compared to Col-0, we measured viral DNA, RNA and protein levels. Viral DNA accumulation was attenuated for both CaMV strains in *lsm1* and *dcp5*, but not in the *xrn4* mutant (Figure 3C). Viral DNA is produced through reverse transcription of the viral 35s RNA and interestingly, the levels of 35s RNA were only mildly reduced for CM1841 and remained unaffected for Cabb B-JI in *lsm1* and *dcp5* (Figure 3D), suggesting that reduced DNA levels could be caused by defects in viral RNA usage in translation or reverse transcription rather than RNA production. Indeed, a western blot analysis showed that the viral inclusion protein P6, the coat protein P4 and the virion-associated protein P3 accumulated less in both *lsm1* and *dcp5* but not *xrn4* (Figure 3E). This was further confirmed by a direct enzyme-linked immunosorbent assay (ELISA), showing reduced quantities of viral proteins P3 and P6 (Figure 3F). In combination, the impairment of CaMV disease in PB mutants indicates a pro- viral role of PBs during CaMV infection. Virus accumulation was specifically impaired in mutants defective in PB biogenesis and decapping (*lsm1* and *dcp5*), but not in exonucleolytic RNA decay (*xrn4*). Both viral strains were similarly impaired in DNA and protein accumulation, with unaltered to modestly reduced RNA accumulation. Owing to the similarities between the two strains, we continued our subsequent analysis with the milder CM1841 strain.

### CaMV RNA capping, stability and TRV-induced gene silencing are not affected in *lsm1* background

The established role of PBs in RNA decapping and degradation led us to test whether these PB associated functions were acting on viral RNA during infection, as the seemingly unaltered viral RNA levels in *lsm1* and *dcp5* mutants could still be explained by a combination of reduced transcription and a defect in RNA decay. To determine the capping levels of viral RNAs in Col-0 and *lsm1* plants, we performed an RNA-pulldown with cap-specific antibodies (Golisz *et al*., 2013). We found known targets of LSM1-mediated decapping to be more abundant in their capped form in the *lsm1* mutant as expected from previous studies (Golisz *et al*., 2013; Perea-Resa *et al*., 2012), while the capping levels of CaMV 35s and 19s RNA did not differ between Col-0 and *lsm1* (Figure 4A). Furthermore, a comparison of known LSM1 targets between the control and CaMV infected samples showed that viral infection does not influence decapping of those endogenous targets, although we cannot exclude the possibility that other targets might be affected (Figure 4B). Unaltered capping of viral RNA was further supported by a cap-sensitive exonuclease digest of total RNA from infected plants, showing identical susceptibility of viral 35s RNA isolated from the *lsm1* mutant compared to Col-0 (Figure 4C). Considering the possibility of PB mediated decapping-independent RNA decay, we also tested whether the decay rate of viral 35S RNA was altered in *lsm1* mutants by quantifying RNA from infected rosettes in a time course after inducing transcriptional arrest through Cordycepin treatment (Sorenson *et al*., 2018). CaMV RNA was remarkably stable and showed no sign of degradation after 120 min of transcriptional inhibition (Figure 4D, left panel). A longer treatment time of eight hours still showed no evident degradation of viral RNA (supplemental Figure 5A), indicating that the viral RNA is strongly protected. The degradation profile of a known target of LSM1 dependent decapping AT4G32020 (Golisz *et al*., 2013) confirmed the transcriptional inhibition and LSM1 dependent effects (Figure 4D, right panel). Our results support that viral RNAs are not major targets of LSM1-dependent decapping or decay and thus, these dysfunctions in *lsm1* and *dcp5* are not likely to cause the reduced virus accumulation.

**Figure 4:**
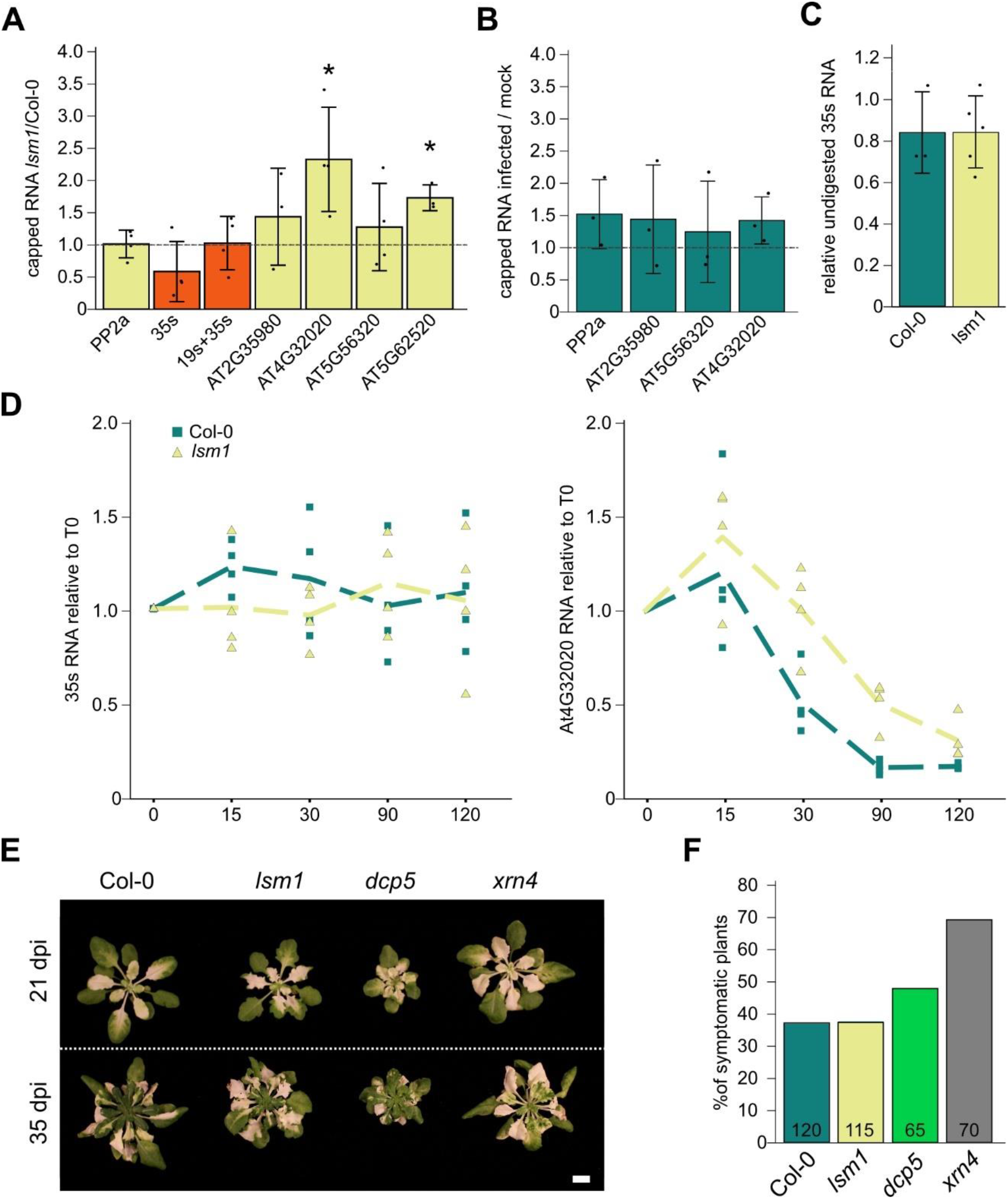
PBs do not regulate viral RNA stability. (A) RNA levels detected after cap-dependent pulldown in infected *lsm1* compared to Col-0 plants for the housekeeping gene *PP2a*, viral RNA and four LSM1 targets. Bars represent mean from independent pulldowns from independent infections (n = 4). (B) RNA levels detected after cap-dependent pulldown on endogenous RNAs from CaMV infected tissue compared to mock infected. Bars represent mean from independent pulldowns from independent infections (n = 3). (C) Amount of viral 35s RNA in Col-0 and *lsm1* mutant detected after 1h of XRN1 treatment. Bars represent the mean from independent digestions from independent infections (n = 3 for Col-0; n = 5 for *lsm1*). Statistical significance was determined by Student’s t test for A-C (*P < 0.05). (D) Transcript decay profiles for viral *35s* and *AT4G32020* RNA after transcriptional arrest. Dotted line represents average of four biological replicates, single experiments are shown by circles (Col-0) and triangles (*lsm1*). Sampling timepoints are indicated on x-axis (0-120 min past treatment). (E) Representative images of VIGS phenotype in indicated genotypes at 21 and 35 dpi with TRV-PDS (scalebar = 1 cm). (F) The percentage of TRV-PDS infected plants that showed whitening at 21dpi. Numbers in bars indicate total number of plants scored.

RQC mutants, are generally prone to initiate RNA silencing against highly expressed RNAs such as transgenes and viral RNAs (Liu & Chen, 2016). Virus-induced gene-silencing (VIGS) is commonly used to silence endogenous genes by including sequence homology into viral genomes, and in the tobacco rattle virus-*Phytoene Desaturase* (TRV-PDS) system infection results in whitening of leaves upon *PDS* silencing (Ratcliff *et al*., 2001). When using this system to compare the general efficiency of VIGS in Col-0, *lsm1, dcp5* and *xrn4* plants, we could not detect increased whitening, delayed recovery, or a higher number of symptomatic plants for *lsm1* and *dcp5*, while *xrn4* showed a clearly enhanced VIGS phenotype (Figure 4E and 4F). Previously, both *xrn4* and a hypomorphic DCP2 mutant *increased transgene silencing 1* (*its1*) were shown to have enhanced VIGS (Ma *et al*., 2019), together indicating that at least TRV- initiated VIGS does not rely similarly on LSM1 and DCP5 as on XRN4 or DCP2. In addition, we tested the expression levels of five target genes of RDR6-dependent *trans*-acting siRNAs (tasiRNA), four of which were shown to be de-repressed during CaMV infection through RNA silencing suppression by the viral P6 protein (Shivaprasad *et al*., 2008). In agreement with their earlier results, we detected a de-repression of *TAS1* and *TAS2*, but not *TAS3* targets during CaMV infection and this de-repression was consistent in *lsm1* and *dcp5* mutants, except for a significantly lower expression of AT1G62590 in *dcp5* (supplemental Figure 5B). Importantly, comparable de-repression of tasiRNA targets by CaMV supports that P6-mediated suppression of RNA silencing is functional in *lsm1* and *dcp5* mutants. Altogether, these results suggests that the canonical PB functions RNA decapping and degradation are not acting on the viral RNA to a detectable extent and thus do not explain the discrepancy between viral RNA and DNA / protein levels in *lsm1* and *dcp5* mutants. Furthermore, *lsm1* and *dcp5* do not show elevated levels of VIGS, distinguishing them from the PB mutants *its1* and *xrn4* (Ma *et al*., 2019) and CaMV P6 is still capable of suppressing RDR6-mediated tasiRNA-dependent silencing on endogenous targets in these mutants.

### LSM1 and DCP5 enhance translation of viral RNA

After establishing that the dynamics of viral RNA levels in the mutant backgrounds behave indistinguishable from wild type, we speculated that not the integrity of the RNA itself, but the translation of it is impaired in *lsm1* and *dcp5*. To test this hypothesis, we performed polysomal profiling on CaMV infected Col-0, *lsm1* and *dcp5* plants. Notably, CaMV infected tissue had increased polysomes compared to mock, and this increase was equally observed in the mutants (Figure 5A), ruling out any global defect in translation. The polysome profiles were divided into twelve fractions (F1-F12) ranging from monosomes to heavy polysomes, followed by RNA extraction for quantification. While all previous assays revealed none to minute differences, the dramatic reduction of viral RNA at polysomes was striking in both *lsm1* and *dcp5*, and specific when compared to the unaltered patterns of the control *PP2a* (Figure 5B and 5D). Importantly, viral RNA and *PP2a* levels were comparable between Col-0 and the mutants in the input lysates used for polysome fractionation (Figure 5C). Considering that both mutants are known to affect P-body formation and show a similar defect in viral RNA translation, we believe that higher-order PB complexes rather than their single proteins are important in the process. Based on these results we propose that PB components promote translation of viral RNA, resulting in enhanced production of viral proteins, DNA and particles.

**Figure 5:**
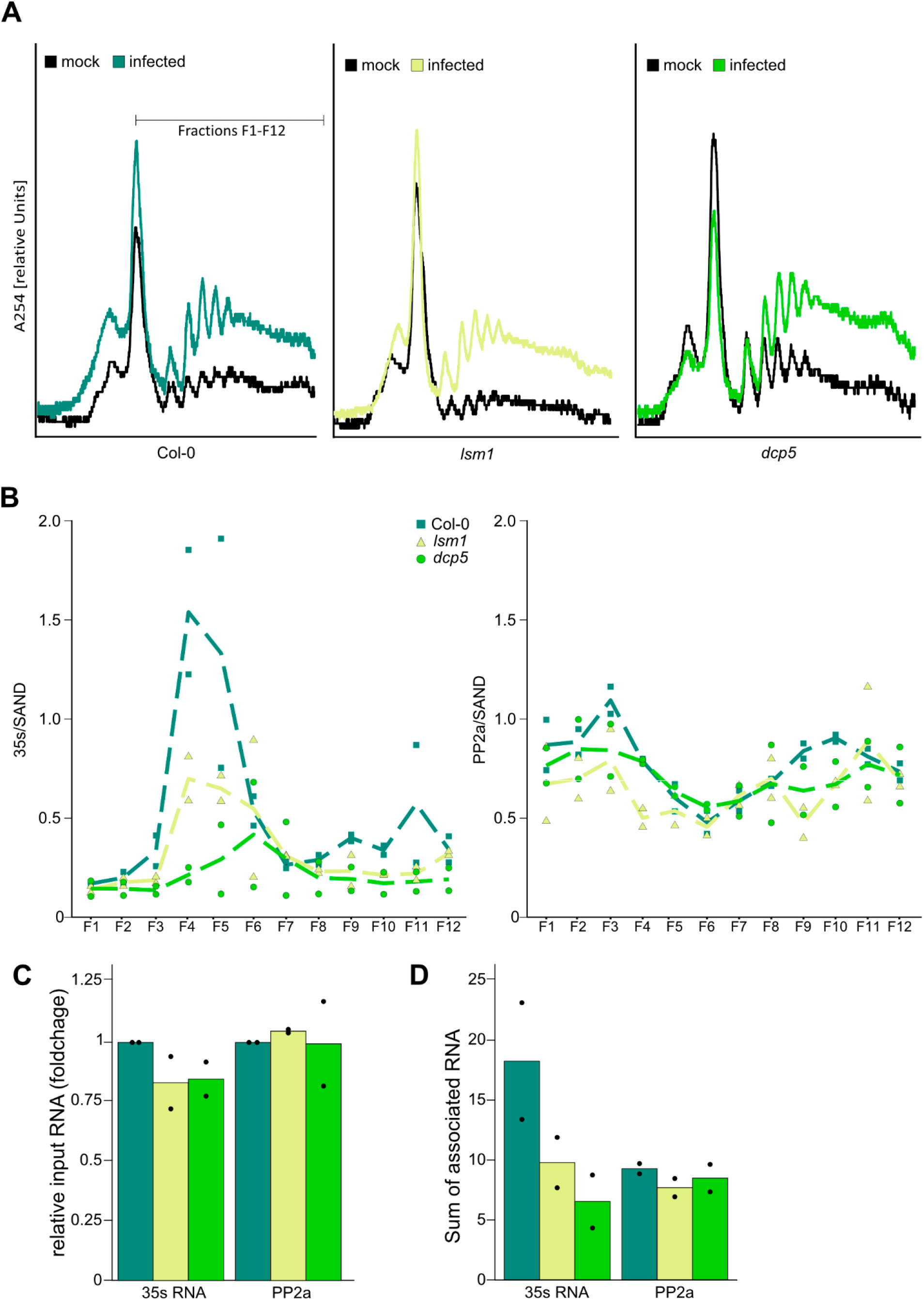
Polysome-association of viral RNA depends on PB components. (A) Polysome Profiles of Col-0, *lsm1* and *dcp5* at 21 dpi CaMV infection. 12 fractions (F1-F12) were taken for polysome associated RNA analysis. (B) RNA abundance in polysome fractions F1-F12 measured for viral *35s* RNA and *PP2a*. Experiment was performed two times using timely separated independent infections. Fractionated RNA was normalized to input RNA and the housekeeping gene *SAND*. Dotted lines represent average of biological replicates, squares (Col-0), triangles (*lsm1*) and circles (*dcp5*) represent single experiments. (C) Relative RNA (foldchange) in input lysates for *35s* RNA and *PP2a* normalized to *SAND*. Dots represent single experiments. (D) Sum of Ribo-Polysome associated RNA in indicated genotypes from (B). Dots represent single experiments.

### Defects in PB integrity expose CAMV to RNA silencing but not NMD

Since PBs are at the heart of RNA triage and a hub for major RNA surveillance mechanisms, we tested whether the disruption of LSM1 and DCP5 exposes CaMV to translational repression via NMD or RNA silencing. To this end, we characterized viral infections in combinatorial mutants of these pathways. CaMV titers were not affected in the previously described NMD mutant *upf1- 5*, although the plants showed a higher relative fresh weight compared to Col-0 (Figure 6A-C). Interestingly, the double mutant *dcp5/upf1* caused the same titer defect as *dcp5* single mutant, showing that this reduction is NMD independent. A previous study found that overexpression of CaMV P6 protein led to a relieve of suppression on several NMD targets containing different NMD marks, including premature termination codons (PTC) and long upstream ORFs (uORFs) (Lukhovitskaya & Ryabova, 2019). During CaMV infection however, we could only detect de-repression of PTC-carrying targets *SMG7* and *RPS6*, suggesting that CaMV specifically represses this NMD mechanism (Figure 6D), possibly to protect against the numerous PTCs present in polycistronic viral RNA. A comparison of transcript levels in infected tissues between Col-0, *dcp5* and *upf1* revealed that NMD target transcription profiles in *dcp5* are more similar to Col-0 than *upf1*, uncoupling NMD regulation during CaMV infections from DCP5 functions (Figure 6E). Polysome association was specifically enriched on the PTC-carrying targets in *dcp5*, showing that DCP5 acts in translational repression on these endogenous targets and that viral RNA is regulated differently from these canonical NMD targets (Figure 6F).

**Figure 6:**
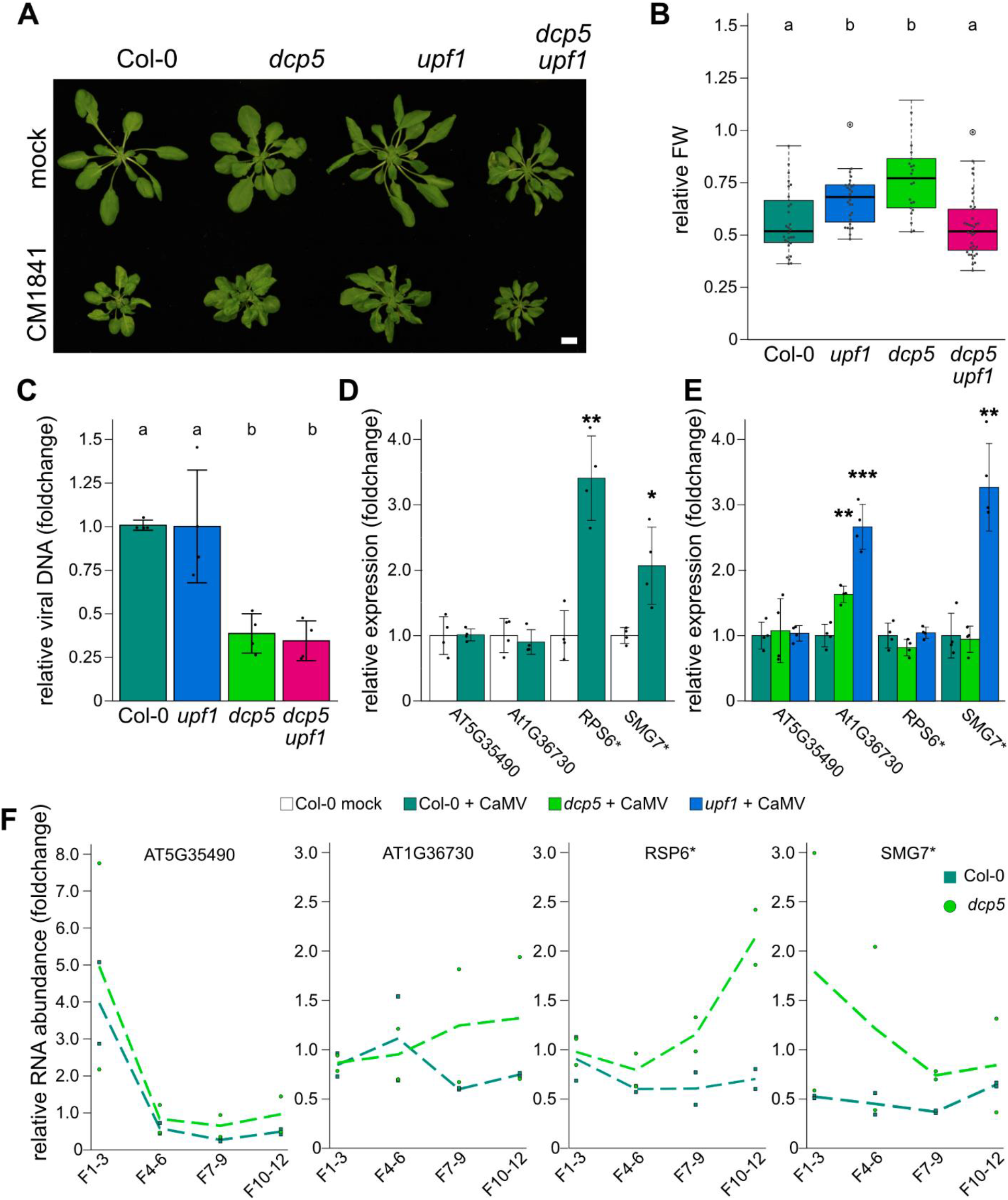
Attenuation of CaMV titers in *dcp5* mutants is not mediated through NMD. (A) Infection phenotype of indicated genotypes at 21 dpi with CM1841 (lower panel) and mock (upper panel) (scale bar = 1 cm). (B) Relative fresh weight of infected / control plants (n=25-30). Solid lines represent median of the data, diamonds the average. Letters indicate statistical groups determined by Kruskal-Wallis test, followed by Wilcoxon sum rank test (alpha = 0.05). (C) Viral DNA accumulation in systemic leaves at 21 dpi, determined by qPCR. Values represent means ± SD (n = 4) relative to Col-0 plants and normalized to *18s* ribosomal DNA as the internal reference. (D) Relative expression of NMD targets in mock and CM1841 infected rosettes 21 dpi determined by qRT-PCR. Values represent means ± SD (n = 4) relative to Col-0 plants and normalized to *PP2a* as the internal reference. * indicates two PTC-containing targets *RSP6* and *SMG7*. (E) Relative expression of NMD targets in CM1841 infected Col-0, *dcp5* and *upf1* plants determined by qRT-PCR. Values represent means ± SD (n = 4) relative to infected Col-0 plants and normalized to *PP2a* as the internal reference. Statistical significance was calculated by two-sided student t-tests (* >0.05; **>0.01; ***>0.001) for C-E. (F) Polysome-associated RNA profiles of indicated genes. Fractions (F) corresponding to fractions in Fig5A. Experiment was repeated twice independently and are the same as in (Figure 5B). RNA was normalized to input RNA and the housekeeping gene *SAND*. Dotted lines represent average of biological replicates, squares (Col-0) and circles (*dcp5*) represent single experiments.

To test whether the translation repression of viral RNA in PB mutants is mediated through the RNA silencing machinery, we crossed *lsm1*, *dcp5* and *xrn4* mutants with *rdr2*, *rdr6*, *dcl2/dcl3/dcl4 (dcl234)* plants (Deleris *et al*., 2006; Allen *et al*., 2004; Xie *et al*., 2004). Higher order mutants, as well as their parental lines were infected with CM1841 and virus disease analyzed at 21 dpi. The mutant alleles tested for *rdr2*, *rdr6* and *dcl234* did not affect plant growth and exhibited a fresh weight loss comparable to Col-0 during CM1841 infection. It is noteworthy that the higher order mutant phenotypes were most similar to the parental PB mutant, showing that the developmental phenotype of *lsm1* and *dcp5* is not alleviated through disruption of RNA silencing (Figure 7A-B). While viral titers remained at Col-0 level in *rdr2*, *rdr6* and *dcl234*, disrupting RDR6, as well as DCL functions rescued the viral titer defect in *lsm1* and *dcp5* backgrounds (Figure 7C). Knock-out of nuclear RNA polymerase RDR2 that is mainly implicated in transcriptional gene silencing (TGS) did not restore CaMV titers (Figure 7C). A protein blot for the viral P6 protein confirmed restoration of viral infection in combinatorial mutants (Figure 7D). Combinations of *xrn4* with either *rdr2* or *rdr6*, respectively, did not alter viral titers at 21 dpi (supplemental Figure 6A-B). It is, however, noteworthy that while viral 35s RNA levels were not affected in any other genotype (supplemental Figure 6C), *xrn4/rdr6* exhibited a slight, but consistent elevation of 35s RNA, suggesting that both pathways can restrict viral RNA accumulation in a compensatory manner. Together, our results show that defect of viral protein production in *lsm1* and *dcp5* mutants is mediated through the cytoplasmic PTGS pathway governed by RDR6 and DCL proteins, and not the NMD pathway. This establishes these PB components as direct antagonists to RNA silencing during CaMV infection and as an escape route for the virus to circumvent targeting by antiviral silencing.

**Figure 7:**
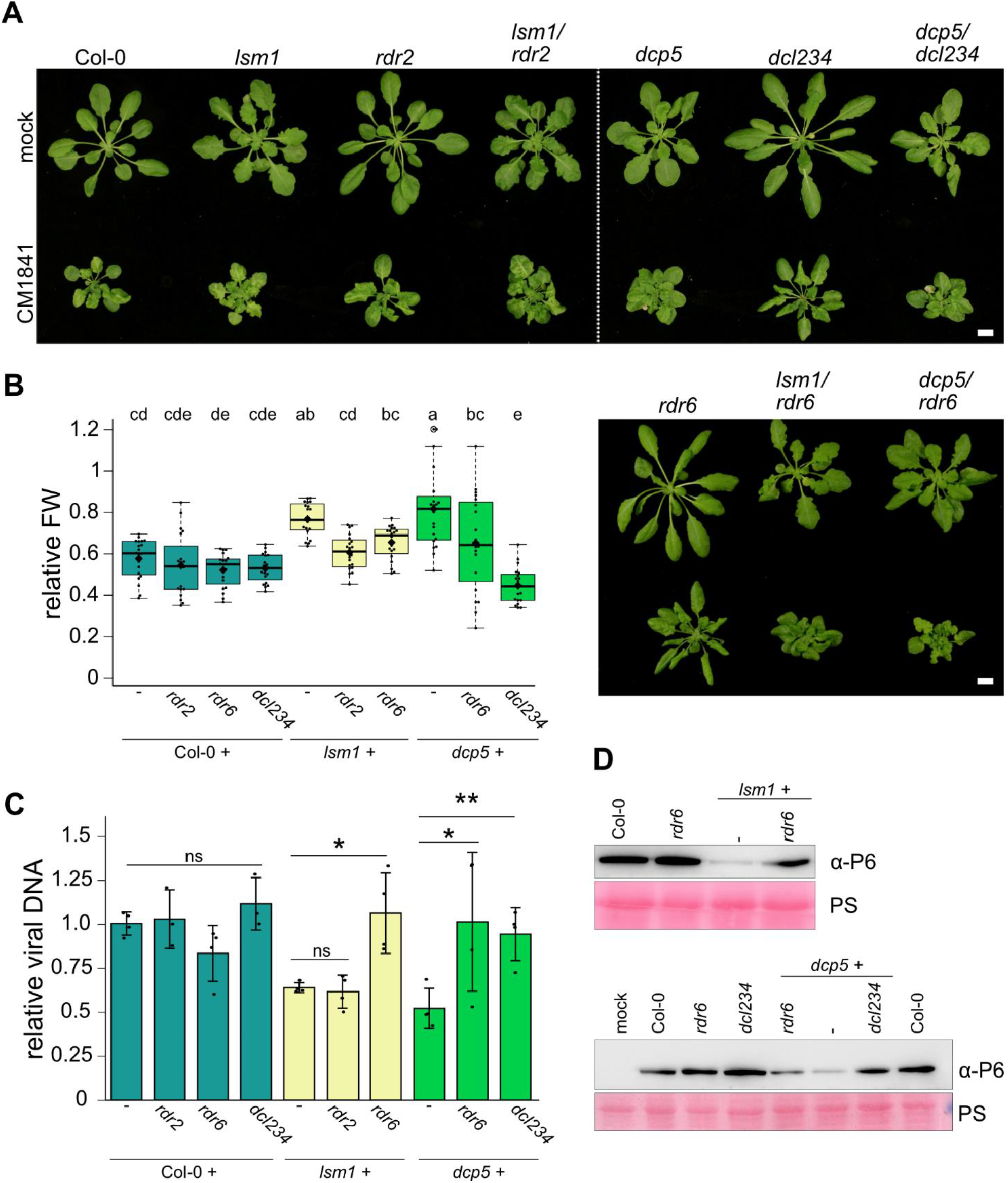
Impairing RDR6-mediated RNA silencing in PB mutants restores CaMV infection. (A) Infection phenotype of indicated genotypes at 21 dpi with CM1841 (lower row) compared to control (upper row). (Scale bar = 1 cm). (B) Relative fresh weight of infected / control plants (n=20). Solid lines represent median of the data, diamonds the average. Letters indicate statistical groups determined by one-way ANOVA followed by Tuckey HSD test (alpha = 0.05) (C) Viral DNA accumulation in systemic leaves of Col-0 and mutant plants at 21 dpi, determined by qPCR. Values represent means ± SD (n = 4) relative to Col-0 plants and normalized to *18s* ribosomal DNA as the internal reference. Statistical significance was calculated by two-sided student t-tests (* >0.05; **>0.01; ***>0.001). Infections experiments are replicated three times independently. (D) Immunoblot analysis of CaMV P6 protein in systemic leaves of indicated genotypes. Total protein samples were extracted at 21 dpi and probed with anti-P6. Ponceau S (PS) staining served as loading control.

## DISCUSSION

### Co-option of PB proteins aids CaMV infection

In contrast to animal virus infections that often trigger a ubiquitous shutdown of translation as part of the hosts antiviral defense (Walsh *et al*., 2013), most plant virus infections have little effect on global *in planta* translation patterns (Li *et al*., 2019; Meteignier *et al*., 2016; Ma *et al*., 2015). CaMV is exceptional in causing a substantial increase of polyribosomes indicative of hyperactivated translation in turnips (Park *et al*., 2001) and Arabidopsis (this study). Translation of CaMV’s polycistronic 35s RNA is a complex process, including mechanisms of leaky scanning and transactivation (Pooggin & Ryabova, 2018). The viral transactivation factor P6 is essential for the translation of downstream ORFs on the 35s RNA (Bonneville *et al*., 1989) through its interaction with a multitude of translation-associated proteins, including the translation initiation factor eIF3g, components of the large ribosomal subunit, the RISP complex and the TOR kinase (Schepetilnikov *et al*., 2011; Park *et al*., 2004).

In the current study, we identified two PB components as important factors for CaMV infection. Traditionally, PB components function as translational repressors for endogenous RNAs (Jang *et al*., 2019; Xu & Chua, 2009; Brodersen *et al*., 2008). Yet, it is possible that viruses have evolved to exploit this pathway for translational targeting of their RNAs as we show for CaMV, where PB components promote viral RNA translation, possibly through their co-assembly in viral factories to aid with ribosome association and escape RNA silencing.

When using four established PB marker proteins, we found distinct localization patterns in non-stressed conditions, and a co-assembly of VCS, LSM1a, DCP5 and DCP1 into granules after heat stress. Our results thus support that PBs containing the higher-order decapping-complex are stress-induced in accordance with previous findings (Perea-Resa *et al*., 2016; Motomura *et al*., 2015; Xu & Chua, 2012), while the constitutive microscopic foci of DCP1, DCP5 and VCS are unlikely to have a high decapping activity and may serve other functions including storage of translationally repressed RNAs (Courel *et al*., 2019; Hubstenberger *et al*., 2017). During CaMV infection, three of the four tested PB components localized to viral factories (VFs). These VFs are structurally different from the membranous RNA virus replication complexes (den Boon & Ahlquist, 2010), and are mainly composed of viral protein P6 and RNA (Schoelz & Leisner, 2017; Martelli & Russo, 1977). This makes CaMV VFs similar to eukaryotic RNA granules, differing from other membranous viral replication complexes, yet they still serve the same purpose to compartmentalize and shield virus replication processes from cellular defenses and optimize particle production. Considering the presence of several PB components in CaMV VFs, we initially hypothesized that RNA granule proteins are either co-opted by the virus to aid infection or targeted to VFs for antiviral defense including viral RNA decay. PB-dependent mRNA degradation and translational repression is selective (Jang *et al*., 2019; Sorenson *et al*., 2018; Hubstenberger *et al*., 2017; Tani *et al*., 2012; Xu & Chua, 2009) and since the viral RNA was stable independent of LSM1-mediated de-capping, these canonical PB functions are unlikely to be associated with VFs. Instead, our data shows that PB components aid CaMV infection as virus accumulation is severely reduced in *lsm1* and *dcp5* mutants. This establishes PBs as pro- viral components in CaMV infection, similarly as observed for other plant viruses (Zhang *et al*., 2020; Hafrén *et al*., 2015; Ye *et al*., 2015). However, as PBs were found to limit viral infections in other contexts (Ng *et al*., 2020; Li & Wang, 2018), it appears that viruses have evolved individually to cope with the PB associated functions that also serve in general plant innate immunity (Chantarachot *et al*., 2020). An intriguing question is how PB proteins are recruited to VFs, and the exclusion of DCP1 suggests specificity to the process. Currently, we consider two possible, and not mutually exclusive, mechanisms associating PBs and VFs (Figure 8). First, a direct interaction of PB proteins with viral proteins, as is suggested by the viral P6 protein binding to the PB scaffold protein VCS in yeast (Lukhovitskaya & Ryabova, 2019). Second, the binding of PB components to viral RNA. CaMV produces its viral RNAs in the nucleus utilizing the canonical transcription machinery of the host (Schoelz & Leisner, 2017), but it is not clear what mechanisms regulate targeting of viral RNAs to VFs and furthermore its partitioning between translation and replication-associated reverse transcription. A PB-dependent RNA shuttling mechanism is a possibility that could be more commonly co-opted by viruses as Lsm1p facilitates *Brome mosaic virus* transport to replication factories in yeast (Díez *et al*., 2000).

**FIG 8:**
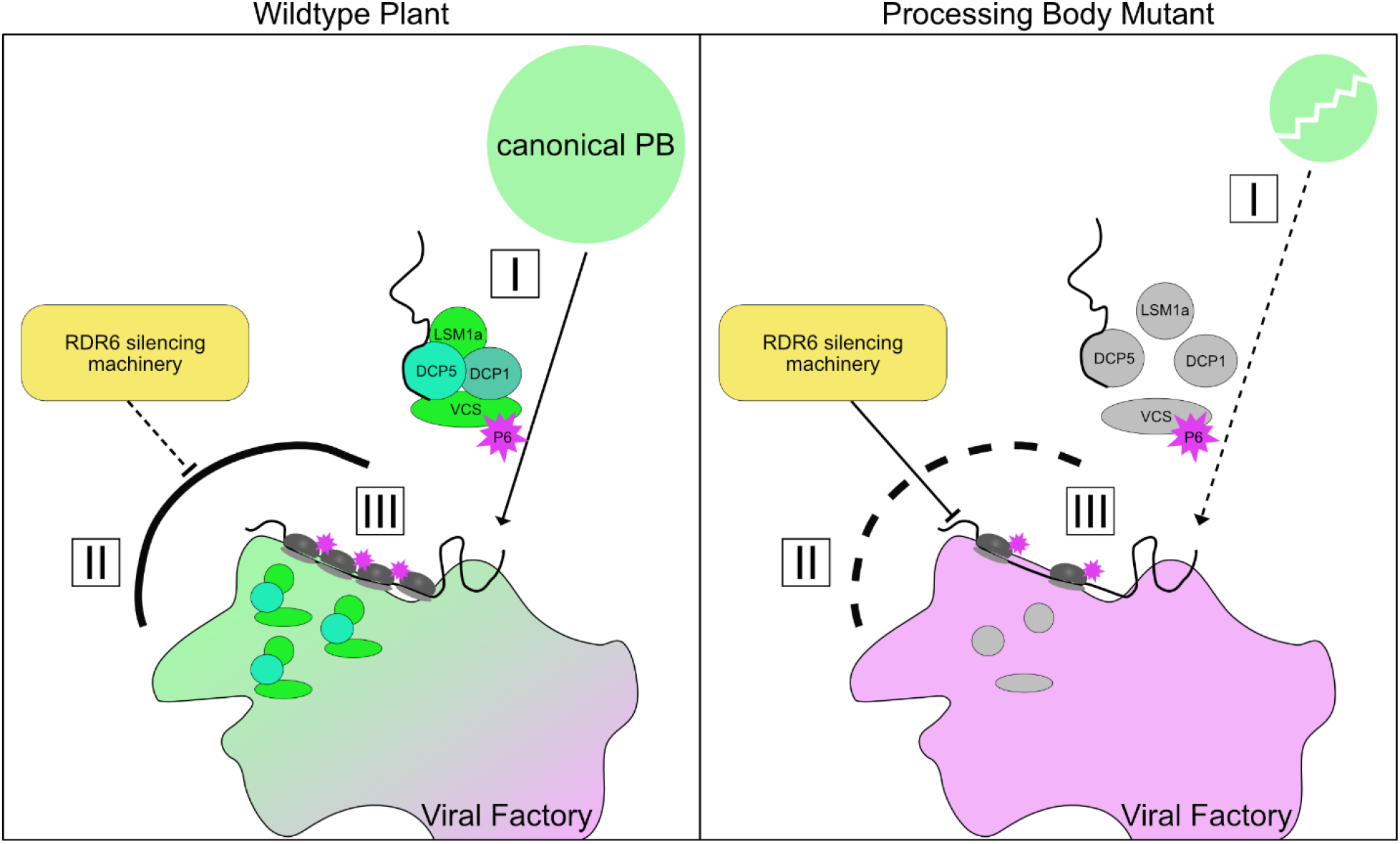
Proposed model of translational repression on viral RNA in PB mutants In wildtype plants (left side): PB components are I) co-opted into viral factories in complexes through viral RNA binding, VCS-P6 binding and/or other unknown mechanisms. In the viral factories, PB proteins II) shield viral RNA from antiviral RDR6-mediated RNA silencing, and by this directly or indirectly III) facilitate translation of viral RNAs and enhance virus accumulation. In PB mutants (right side) the VF-targeted complex is broken, leading to less efficient viral RNA regulation. In turn, viral RNA becomes susceptible to RNA silencing mediated translational repression, resulting in decreased viral accumulation.

### Viral RNA is regulated differently than canonical NMD targets

The polycistronic viral 35s RNA contains several potential triggers for RNA surveillance mechanisms, including premature stop codons (PTCs), a large stem-loop and extremely high expression levels. PTCs are key signatures for NMD (Peltz *et al*., 1993) and in plants, this pathway was shown to suppress infections of PTC carrying RNA viruses (Garcia *et al*., 2014). A primary reason for addressing the NMD regulator UPF1 in this study is its largely shared protein interactome with DCP5 (Chicois *et al*., 2018) and general coupling with PBs (Raxwal *et al*., 2020; Lejeune *et al*., 2003). However, CaMV showed UPF1-independent accumulation in both *upf1* and *dcp5 upf1* mutants, disconnecting the NMD pathway from the pro-viral function of DCP5. Furthermore, we also found that endogenous targets of NMD decay were stabilized during infection in a DCP5-independent manner, suggesting that CaMV suppresses mRNA decay occurring through the NMD pathway irrespective of these PB components. Interestingly, viral P6 protein was recently shown to suppress NMD involving a possible interaction with the PB component VCS (Lukhovitskaya & Ryabova, 2019). More generally, we speculate that NMD inhibition during CaMV infection is a consequence of increased translation re-initiation known to represses PTC-triggered NMD (Karousis & Mühlemann, 2019; Raimondeau *et al*., 2018). Finally, we found that endogenous PTC transcripts stabilized by CaMV, were more associated with polysomes in *dcp5*, suggesting that PBs can mediate translational repression of NMD targets. This finding supports our general understanding of NMD and PB co-operation in removing unwanted transcripts from translation, and the proposed model of P6 association with the PB component VCS to suppress PTC-triggered NMD. Nevertheless, the opposite behavior of viral RNA means that other, more decisive, mechanisms override and drive its translational repression in *dcp5* and *lsm1*.

### CaMV utilizes PB proteins to enhance protein production and evade RNA silencing

We show that RNA silencing components RDR6 and DCLs are necessary for the DNA and protein defect occurring in PB mutants. Viruses have frequently evolved to evade and suppress the main plant antiviral pathway RNA silencing, and disruption of it can lead to hyper-infections (Csorba *et al*., 2015; Garcia *et al*., 2014; Garcia-Ruiz *et al*., 2010; Morel *et al*., 2002). However, CaMV is resilient to RNA silencing (Blevins *et al*., 2011) and we show that the viral escape strategy includes PB components for maintenance of efficient viral translation in the presence of cytoplasmic RNA silencing. We did not observe an activation of silencing using the TRV VIGS system in *lsm1* and *dcp5*, and importantly, we also found comparable de-repression of tasiRNA targets in PB mutants, suggesting that the P6-mediated suppression of the DRB4/DCL4 node of PTGS (Haas *et al*., 2008) is not attenuated. While it is not possible to rule out that either RNA silencing is hyper activated in a manner not detected through VIGS or P6 suppression is weaker in PB mutants, we favor the hypothesis that PB dysfunctions expose the virus to RNA silencing mechanisms that it has not adapted to and normally avoids (Figure 8). Notably, a primary link between RNA silencing and PBs in plants was established from forward genetic screens of induced transgene silencing, identifying both *xrn4* (Gazzani *et al*., 2004) and *dcp2* (Thran *et al*., 2012). Analogous to compromised CaMV infection in *lsm1* and *dcp5*, transgene silencing in *xrn4* and *dcp2* mutants depended on RDR6. However, a fundamental difference is that the latter involved the degradation of the silenced transcripts, while CaMV silencing acts on translation efficiency but not RNA degradation. Translational repression through ribosome stalling especially during stress adaption has recently emerged as a major function of RNA silencing (Kim *et al*., 2021; Iwakawa *et al*., 2020; Wu *et al*., 2020). Combining the concept of substrate channeling and competition between PBs and siRNA bodies (Jouannet *et al*., 2012) with a viral infection context, we suspect that the association of PB components with CaMV VFs reduces viral RNA template acquisition by the RDR6 silencing machinery. Taken together, we propose that CaMV co-opts PB components to shield it from RNA silencing and evade translational repression. (Figure 8).

## Material & Methods

### Plant Material and Growth Conditions

All mutants used in this study were in the *Arabidopsis thaliana* accession Columbia (Col-0) which was taken as control for all experiments (supplemental Dataset1 Table S1 and S3). Arabidopsis and *Nicotiana benthamiana* plants were grown in walk-in chambers in standard long day conditions (16h light / 8h dark cycle) at 22°C and 65% relative humidity for crossing, propagation, and transient expression assays. For infection experiments, plants were grown in short day conditions (120 mE, 10h light / 14h dark cycle) at 22°C and 65% relative humidity.

### Plasmid Construction, Generation of Transgenic Lines and Transient Expression

The pENTRY clone containing full length Cabb B-JI P6 coding sequence (Hafren *et al*., 2017) was cloned into pGWB654 or pGWB554 vectors under the control of the 35s promoter (Nakagawa *et al*., 2007). Expressor lines were generated for this study by floral dipping (Clough & Bent, 1998), all lines and constructs are listed in supplemental Dataset1 Table S1. Coding sequences of DCP1, DCP5, LSM1a and VCS were amplified from Col-0 plants (primers listed in supplemental dataset1 Table S4), cloned into pENTR/D-TOPO and subsequently recombined in pUBC-dest vector system (pUBN-dest for VCS) (Grefen *et al*., 2010). For transient expression, *Nicotiana benthamiana* leaves were infiltrated with resuspended Agrobacteria (OD 0.2, 10mM MgCl2, 10mM MES pH 5.6, 150µM Acetosyringone) and the constructs analyzed after 48h.

### Virus Inoculation and Quantification

Arabidopsis plants were infected with CaMV or TRV 18 days after germination. The first true leaves were either infiltrated with *Agrobacterium tumefaciens* strain C58C1 carrying CaMV strain CM1841, or TRV RNA1 and 2 (OD 0.15) or mechanically rubbed with turnip-purified particles of Cabb B-JI resuspended in carborundum supplemented phosphate buffer (Martinière *et al*., 2009). Rosettes were harvested 21 dpi in 4 biological replicates, each containing two to three individual plants from which inoculated leaves were removed. For CaMV DNA quantification, 100mg pulverized frozen leaf material was resuspended in 300 µl 100mM Tris buffer (pH 7.5), supplemented with 2% SDS and treated with Proteinase K. Total DNA was precipitated with isopropanol 1:1 (v:v) and viral DNA levels were determined by qPCR and normalized to *18s* ribosomal DNA {Hafren, 2017 #62}. RNA extraction from rosette tissue was performed with the Qiagen RNeasy kit according to manufacturer’s protocol. 500 ng of total RNA were used for first-strand cDNA synthesis with the Maxima First Strand cDNA Synthesis Kit (Thermo Fisher Scientific). qRT-PCR analysis was performed with Maxima SYBR Green/ Fluorescein qRT-PCR Master Mix (Thermo Fisher Scientific) using the CFX Connect Real-Time PCR detection system (Bio-Rad) with gene-specific primers (supplemental Dataset1 Table S3). Viral transcripts were normalized to *PP2a* (AT1G69960) and expression levels determined as described by (Livak & Schmittgen, 2001). Viral proteins were extracted from frozen rosette tissue in 100 mM Tris buffer (pH 7.5), supplemented with 2% SDS. Samples were incubated at 95°C for 5 min in 1x Laemmli sample buffer and cleared by centrifugation. The protein extracts were separated by SDS-PAGE, transferred to PVDF membranes (Amersham, GE Healthcare) and blocked with 8% (w:v) skimmed milk in 1x PBS, supplemented with 0.05% Tween 20. Blots were incubated with 1:2000 diluted primary antibodies α-P3, α-P4 or α-P6 before subsequent addition of secondary horseradish peroxidase (HPR)-conjugated antibodies (1:20.000; Amersham, GE Healthcare). The immunoreaction was developed using the ECL Prime kit (Amersham, GE Healthcare) and was detected in the LAS-3000 Luminescent Image Analyzer (Fujifilm). An ELISA was performed for three independent experiments, with 100 mg infected plant material in 1ml (w/v) 8M Urea buffer. Samples were incubated on high-binding ELISA plates for 6h at 37°C before blocking in 5% skimmed milk. Primary antibodies were added 1:500 dilutions overnight and secondary antibodies 1:1000 dilution for 3h at 37°C. Absorbance was measured at 405nm from 30-120 min after addition of Substrate buffer (PNPP, Thermo Fisher).

### Cap-dependent Immunoprecipitation and XRN1 digest

Immunoprecipitation of m^7^G-capped RNA was performed as described by (Golisz *et al*., 2013). Anti-7-methylguanosine (m7G)-Cap mAb (clone 150-15) were purchased from MBL International Corporation. For the exonucleolytic digest, total RNA was extracted from rosettes 21 dpi and incubated at 37°C with 1U XRN1 enzyme (Thermo Fischer) or in reaction buffer (mock) (Roux *et al*., 2015). cDNA synthesis and qRT-PCR were performed as described in the previous chapter. Transcript levels were normalized to *eIF4a* (At3g13920) (Roux *et al*., 2015; Perea-Resa *et al*., 2012).

### RNA half-life measurement

Rosettes of CaMV infected plants (21 dpi) were vacuum infiltrated with 1mM Cordycepin (Sigma-Aldrich) in buffer (1mM PIPES, pH 6.25, 1mM sodium citrate, 1mM KCl, 15mM sucrose) and placed in a damp chamber. Two plants were harvested per sample corresponding to 0, 15, 30, 60, 120 min after transcriptional inhibition. Total RNA was extracted using Trizol reagent followed by cDNA synthesis and qRT-PCR as described in the previous chapter. RNA levels were normalized to *eIF4a* (At3g13920).

### Ribosomal Profiling

Polysome extraction was performed based on (Mustroph *et al*., 2009) with some modifications. Briefly, 1 ml frozen leaf powder was thawed in 8 ml of polysome extraction buffer (200 mM Tris·HCl (pH 8.0), 200 mM KCl, 35 mM MgCl2, 25 mM EGTA, 1 mM DTT, 1 mM phenylmethanesulfonylfluoride, 100 μg/mL cycloheximide, 1% (vol/vol) detergent mix (20% (w/v) Brij-35, 20% (v/v) Triton X-100, 20% (v/v) Ipegal CA630 and 20% Tween 20), 1% (v/v) polyoxyethylene 10 tridecyl ether), resuspended and kept on ice for 10 minutes. The plant debris was removed by centrifuging at 16 000 g for 15 minutes at 4°C in a JA-25.50 rotor and Avanti J- 20 XP centrifuge (Beckman Coulter). The clear supernatant was then gently poured on top of 8 ml sucrose cushion (100 mM Tris·HCl (pH 8.0), 40 mM KCl, 20 mM MgCl2, 5 mM EGTA, 1 mM DTT, 100 μg/mL cycloheximide in 60% sucrose) in a 26 ml polycarbonate tube (Beckman Coulter). After proper balancing, the samples were centrifuged at 35 000 RPM for 18 hours at 4°C in a 70Ti rotor and L8-M ultracentrifuge (Beckman Coulter). The ribosome pellets were then gently washed with RNase-free water and resuspended in 300 μl of resuspension buffer (100 mM Tris·HCl (pH 8.0), 40 mM KCl, 20 mM MgCl2, 100 μg/mL cycloheximide). The resuspended samples were kept on ice for 30 minutes followed by centrifugation at 16 000 g at 4°C to remove any debris. The RNA content was measured for each sample using Qubit BR RNA assay kit (Thermo Fisher Scientific). The resuspended polysome samples were then loaded on 15% to 60% sucrose gradients and centrifuged at 50 000 RPM in a SW55.1 rotor and L8-M ultracentrifuge (Beckman Coulter). The gradient samples were fractionated using an ISCO absorbance detector (model # UA-5, ISCO, Lincoln, NE) to obtain 12 ribosome-containing fractions of approx. 250 µl and RNA was extracted from individual fractions using Trizol reagent. Single fractions were used for cDNA synthesis and qRT-PCR as described in the previous chapter. RNA levels were normalized to *SAND* (At2g28390) in each fraction and depicted as % of total RNA.

### Confocal Microscopy and Treatments

Micrographs from leaf abaxial epidermal cells were taken with the Zeiss LSM 780 microscope. GFP and RFP signals were detected at 488 nm/490– 552 nm and 561 nm/569–652, respectively. Co-visualization was achieved through sequential scanning mode. For HS conditions, leaves were kept in water at 38°C for 30 min (1h for LSM1-GFP) before imaging. Translational inhibition treatment was achieved through infiltration of young leaves with 200 µM CHX (Sigma-Aldrich), followed by an incubation of 1h before imaging. Images were processed with the ZEN black software (Zeiss) and ImageJ version 1.53b. For quantifications, Z-stacks were Brightness increased and a median filter of 2 pixels applied. Stomata were manually deleted from micrographs and a mask generated through thresholding. Foci were counted using the “Analyse Particles” tool.

### Data Analysis and Statistical Methods

Statistical comparisons of two groups were performed by Student’s T-test. One-way ANOVA followed by a post-hoc Tukey HSD test (α = 0.05) was performed with R v4.0.02 and the R-package “agricolae” (Version 1.3-3; https://cran.rproject.org/web/packages/agricolae/index.html).

### Accession Numbers

Sequence data from this article can be found in the EMBL/GenBank data libraries under accession number(s): DCP1 (AT1G08370), LSM1a (AT1G19120), LSM1b (AT3G14080), DCP5 (AT1G26110), XRN4 (AT1G54490), VCS (AT3G13300), RDR2 (AT4G11130), RDR6 (AT3G49500). DCL2 (AT3G03300), DCL3 (AT3G43920), DCL4 (AT5G20320), UPF1 (AT5G47010)

## Supplemental Data

Supplemental Figure 2: PB double marker lines show co-assembly after HS and during CaMV infection

Supplemental Figure 3: Co-expression of PB components with CaMV proteins in *Nicotiana benthamiana*

Supplemental Figure 4: P6 co-localization with PB components in mock conditions

Supplemental Figure 5: Long-term viral RNA stability and RNA silencing suppression

Supplemental Figure 6: CaMV infection in *xrn4* backgrounds and 35s RNA accumulation

Supplemental Dataset1: Primers and Plant lines used in this study

## ACKNOWLEDGMENTS

We thank Dr. Aayushi Shukla and Dr. Heinrich Bente for their valuable input on this manuscript. We thank Joanna Kufel for sharing the *lsm1a/b* double mutant, Cécile Bousquet-Antonelli for the *xrn4* mutant, James Carrington for the *rdr* and *dcl* mutants This study was supported by the Swedish Research Council VR (grant number 2017-05036) for AH. JH and AB were supported by grants from the Knut and Alice Wallenberg Foundation, the Swedish Governmental Agency for Innovation Systems (VINNOVA) and Bio4Energy, a Strategic Research Environment appointed by the Swedish government. DG was funded by an attractivity grant from the NetRNA LabEx, ANR-10-LABX-0036_NETRNA.

## AUTHOR CONTRIBUTIONS

G.H. and A.H. designed the experiments and wrote the manuscript. G.H. conducted the experiments and analyzed the data together with A.H. Polysome analysis was conducted and analyzed together with A.M. and J.H. D.G. provided the *upf1*, *upf1/dpc5* and DCP5-GFP Arabidopsis lines. All authors edited the manuscript and approved the final version.

---

**Figure S1:**
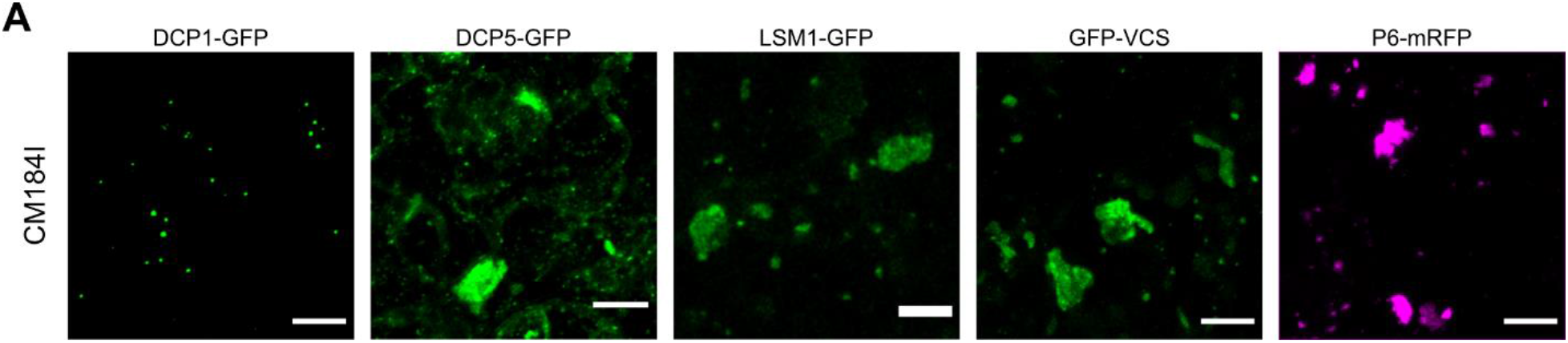
Localization of PB marker proteins in advanced infections. (A) Confocal images composed of confocal Z-stacks of PB markers and CaMV P6 five to seven weeks after infection, imaged genotypes are indicated above micrographs (Scalebar = 10µm).

**Supplemental FIG 2:**
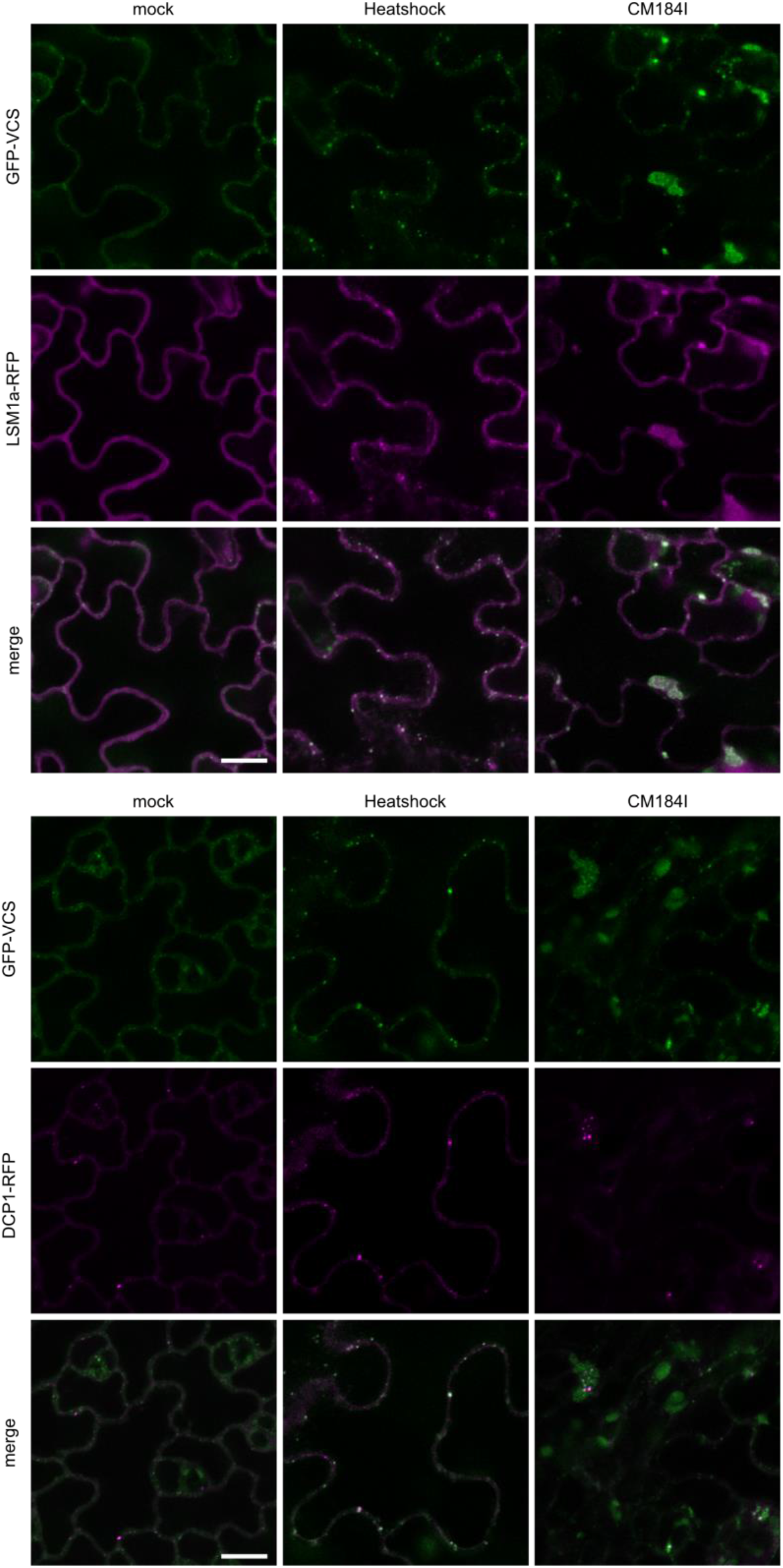
PB double marker lines show co-assembly after HS and during CaMV infection. Co-localization of GFP-VCS with LSM1-RFP (upper panel) and DCP1-RFP (lower panel) in transgenic Arabidopsis 21 days after mock, heat shock or CaMV infection. Images represent single plane micrographs (Scale bars = 10 µm).

**Supplemental Figure S3:**
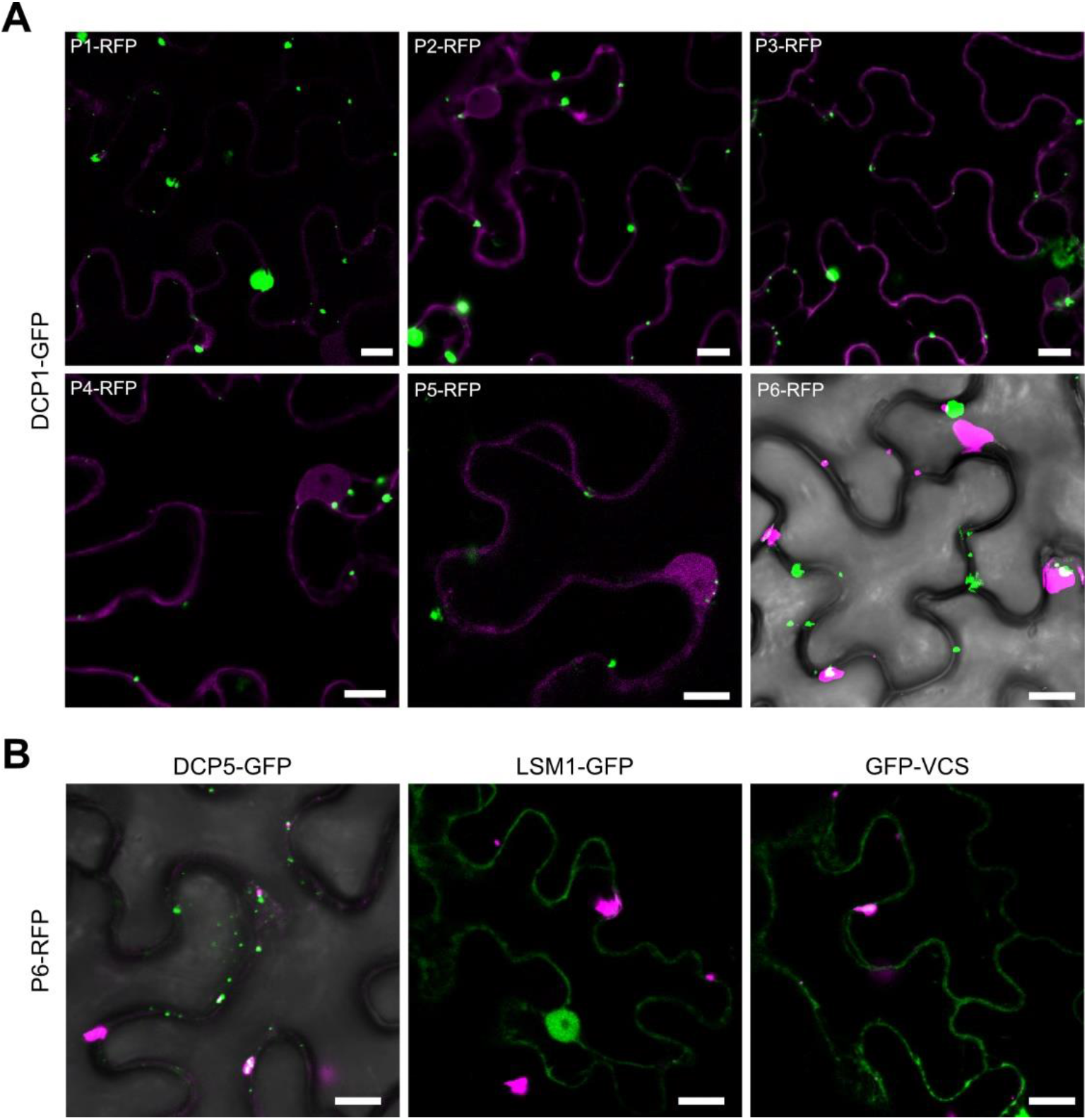
Co-expression of PB components with CaMV proteins in *Nicotiana benthamiana*. (A) Viral proteins fused to tagRFP were co-expressed with AtDCP1-GFP in *Nicotiana benthamiana* leaves. Single plane images were taken 2 days after co-infiltration. Brightfield channel is included when neither of the expressed proteins are soluble enough to outline the cell contour. (Scale bars = 10 µm). (B) PB components were co-expressed with BJI P6-RFP. Single plane images were taken 2 days after co-infiltration in *Nicotiana benthamiana* leaves. (Scale bars = 10 µm). Brightfield channel as in (A).

**Supplemental FIG 4:**
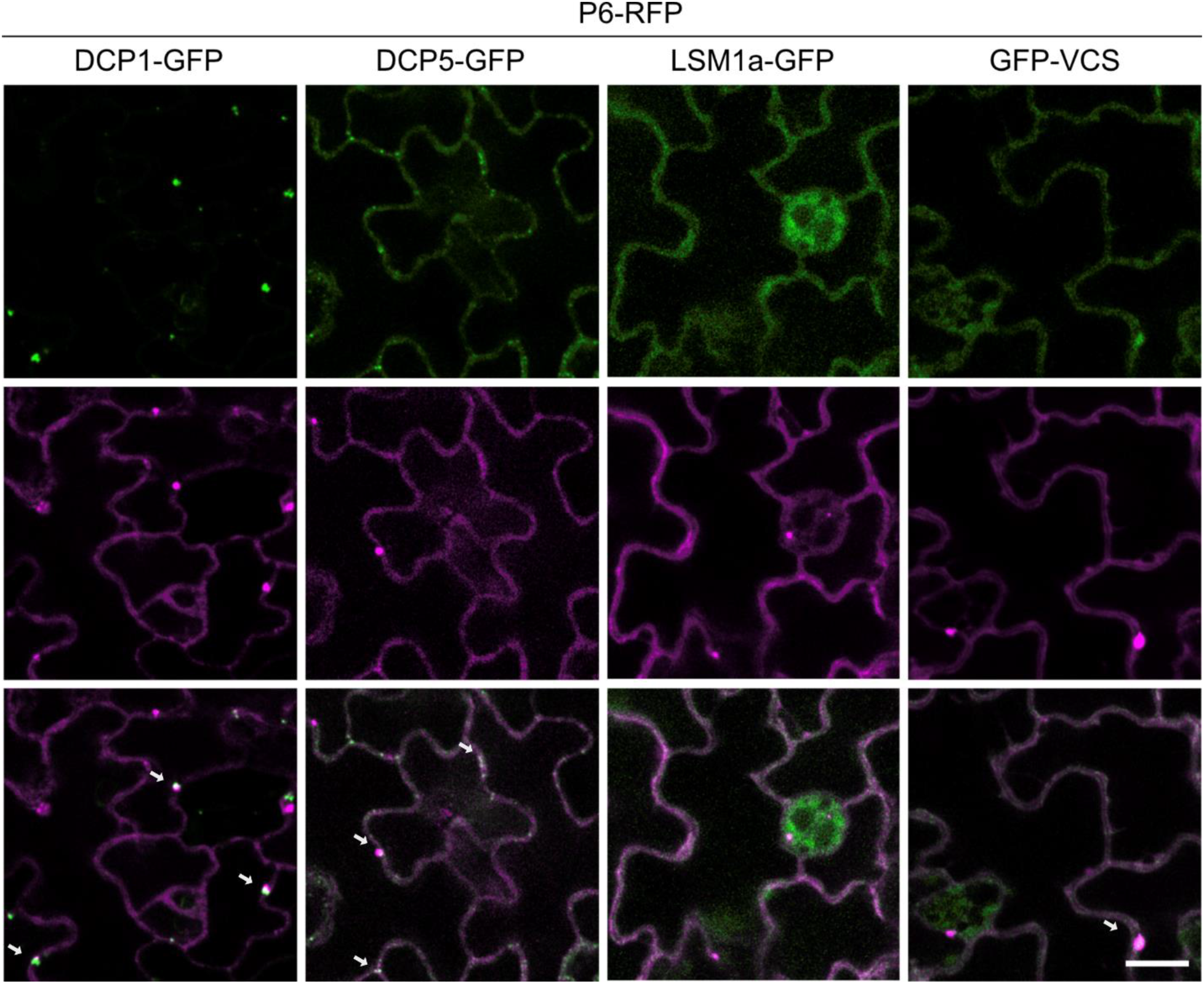
P6 co-localization with PB components in mock conditions. Co-localization of P6-mRFP with PB markers in transgenic Arabidopsis 21 dpi after mock infection. White arrows point to co-localizations. Representative single plane images are shown (Scale bars = 10 µm). Experiments were replicated at least three times with independent transformants.

**Supplementary Figure 5:**
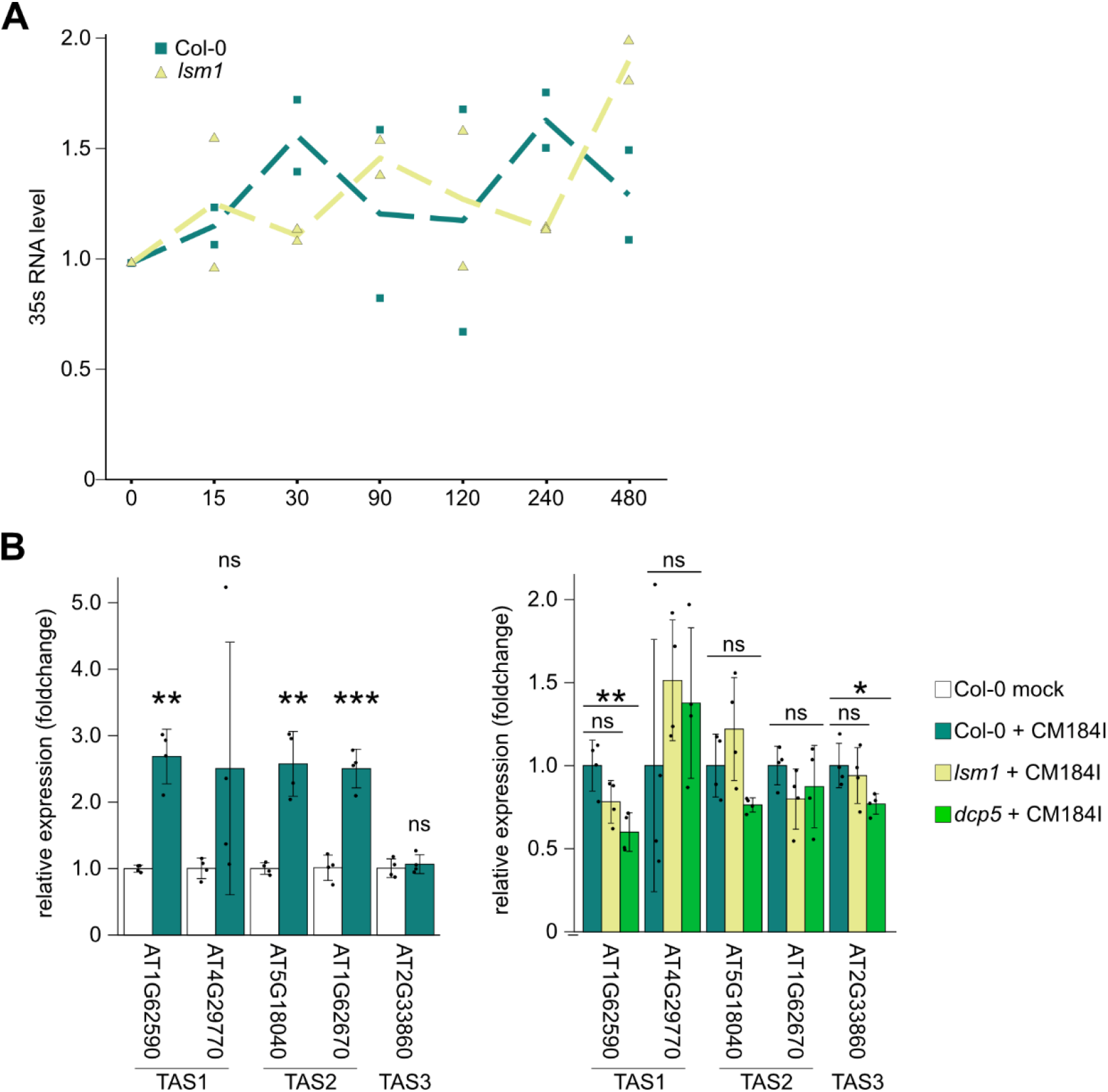
Long-term viral RNA stability and RNA silencing suppression. (A) Long-term transcript decay profiles for viral 35s RNA after transcriptional arrest using cordycepin. Dotted line represents average of two biological replicates, single experiments are shown by circles (Col-0) and triangles (*lsm1*). Sampling timepoints (min after treatment) are indicated on x-axis. (B) Expression of five different tasiRNA targets at 21 dpi with CaMV compared to mock (left side) and compared to Col-0 infection in *lsm1* and *dcp5* (right side) (n=4). tasiRNA generating TAS loci are indicated below the gene identifiers. Statistical significance was calculated by two-sided student t-tests (* >0.05; **>0.01; ***>0.001). Experiments were repeated three times from independent infections.

**Supplemental Figure 6:**
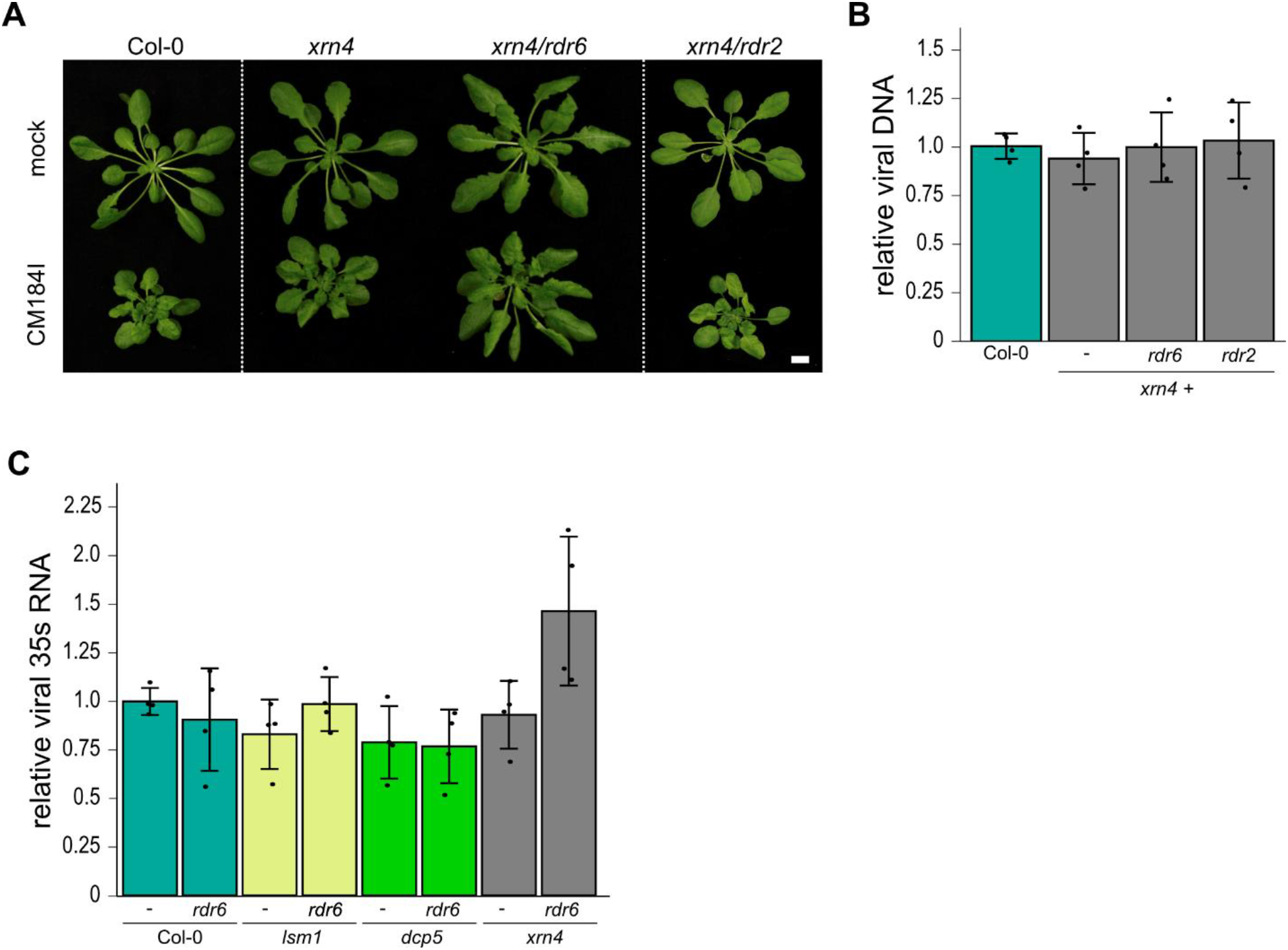
CaMV infection in *xrn4* backgrounds and 35s RNA accumulation. (A) Infection phenotype of indicated genotypes at 21 dpi with CM1841. Scale bar = 1 cm (B) Viral DNA accumulation by qPCR in systemically infected rosettes of indicated genotypes at 21 dpi relative to Col-0 and normalized to *18s* ribosomal DNA for internal reference. Averages from 4 independent infection experiments. (C) Viral 35s RNA accumulation in systemically infected rosettes of indicated genotypes at 21 dpi relative to Col-0 determined by qRT-PCR. Values depict averages from 4 independent infection experiments, relative to Col-0 and normalized to *PP2a*.

